# The yeast CST and Polα/primase complexes act in concert to ensure proper telomere maintenance and protection

**DOI:** 10.1101/2024.12.20.629258

**Authors:** Kimberly Calugaru, Eun Young Yu, Nayim González-Rodríguez, Javier Coloma, Neal F. Lue

## Abstract

Polα/primase, the polymerase that initiates DNA synthesis at replication origins, also completes the task of genome duplication by synthesizing the telomere C-strand under the control of the CST complex. Using cryo-EM structures of the human CST-Polα/primase-DNA complex as guides in conjunction with AlphaFold modeling, we identified structural elements in yeast CST and Polα/primase that promote complex formation. Mutating these structures in *Candida glabrata* Stn1, Ten1, Pri1 and Pri2 abrogated the stimulatory activity of CST on Polα/primase *in vitro,* supporting the functional relevance of the physical contacts in cryo-EM structures as well as the conservation of mechanisms between yeast and humans. Introducing these mutations into *C. glabrata* yielded two distinct groups of mutants. One group exhibited progressive, telomerase-dependent telomere elongation without evidence of DNA damage. The other manifested slow growth, telomere length heterogeneity, ssDNA accumulation and elevated C-circles, which are indicative of telomere deprotection. These telomere deprotection phenotypes are altered or suppressed by mutations in multiple DDR and DNA repair factors. We conclude that in yeast, the telomerase inhibition and telomere protection function previously ascribed to the CST complex are mediated jointly by both CST and Polα/primase, highlighting the critical importance of a replicative DNA polymerase in telomere regulation.

## Introduction

Linear eukaryotic chromosomes are capped at their ends by special nucleoprotein structures called telomeres, which promote genome integrity by stabilizing the ends against aberrant re-arrangements (1,2). Formation of the telomere nucleoprotein complexes is nucleated by numerous copies of a short repeat sequence, which recruit DNA binding proteins and additional factors to execute the end protection function. Aside from end protection, telomeres need to minimize and compensate for DNA loss incurred due to the failure of replication machinery to fully copy the DNA template at chromosome ends, i.e., the end replication problem (3-5). Accordingly, eukaryotic cells have evolved specialized DNA synthesis complexes and regulatory proteins to add telomere DNA onto chromosome ends.

The two polymerases responsible for adding telomere repeats are telomerase and Polα/primase (PP), which sequentially elongate the two strands of telomeres (6,7). First, the telomerase reverse transcriptase uses an embedded segment of a stably associated RNA subunit as template to extend the 3’-end bearing, G-rich strand. The extended G-strand is then converted to double-stranded DNA by PP, a bi-functional enzyme with both primase and DNA polymerase activities (8). Even at telomeres not extended by telomerase, the replication process results in sizable G-strand ssDNA that needs to be converted to duplex DNA (3). PP “fills-in” the telomere C-strand by synthesizing an RNA oligomer using the primase subunits (Pri1 and Pi2) and then extending the RNA using the DNA polymerase (Pol1 and Pol12). Notably, PP is not only essential for telomere maintenance but also for chromosomal replication; it functions as the initiator protein by synthesizing short RNA-DNA oligomers that serve as primers for extension by both the leading and lagging strand polymerases (9). While the enzymatic activities of PP in telomere maintenance and chromosomal replication are identical, they are regulated through different mechanisms. In genome replication, PP is assembled into the replisome through interaction with the CMG complex (10,11), whereas at telomeres, PP is physically and functionally coupled to the CST complex (12-14).

CST (CTC1-STN1-TEN1 in metazoans, plants and some fungi; Cdc13-Stn1-Ten1 in budding yeasts) is an evolutionarily malleable, RPA-like complex that plays critical roles in telomere regulation; it terminates telomerase-mediated G-strand synthesis and promotes C-strand synthesis by stimulating PP. That is, CST prevents excessive telomere elongation and ensures the addition of double-stranded telomere repeats to chromosome ends. Mutations in CST subunits have been shown to trigger aberrant telomere lengthening and/or shortening depending on the context and the nature of the mutations. However, the precise mechanisms of CST in the telomere maintenance pathway remain poorly understood. One unresolved issue is the potential participation of PP in terminating telomerase activity, with some studies indicating that CST alone is sufficient, whilst others supporting the additional requirement for PP (15). Beyond regulating telomere DNA synthesis, the budding yeast CST also plays an essential role in telomere protection; all three subunits are essential and hypomorphic alleles have been shown to trigger DNA damage response and aberrant repair. This additional function of yeast CST may have evolved to compensate for the loss of shelterin proteins – the major protective complexes at telomeres in the great majority of eukaryotes (16).

From the mechanistic perspective, the C-strand synthesis reaction compels interests on its own. Like the PP reaction in chromosomal replication, it undergoes multiple conformational transitions in order to execute 1) RNA synthesis initiation and elongation, 2) the handoff of the RNA from the primase active site to the DNA polymerase activate site, as well as 3) the initiation of DNA synthesis and DNA chain elongation (17). While it is clear that both mammalian and yeast CST can substantially stimulate the synthesis of RNA-DNA products by PP (18,19), whether and how CST modulates the individual steps of this reaction remain poorly understood. Even though the budding yeast *Saccharomyces cerevisiae* has offered many insights on telomere regulation in the past three-plus decades (20), it has presented notable challenges in exploring the molecular mechanisms of CST-PP. Most importantly, the *Sc*CST complex has been refractory to recombinant expression and purification, making it difficult to investigate the CST-PP-mediated reactions *in vitro*. To overcome this challenge, we established an alternative co-expression/purification system for this complex in *C. glabrata* (21). We further purified *Cg*PP and showed that CST stimulates PP by enhancing both the priming and the primase-to-polymerase switch reactions (18).

Herein we report our continued investigation of *Cg*CST-PP. As a point of departure, we leveraged the recent cryo-EM structures of the human CST-PP-DNA pre-initiation complex (PIC), which reveals multiple contacts between CST, PP and the DNA template (22). In combination with AlphaFold modeling and comparative sequence analysis, we identified surface residues in *Cg*Stn1, Ten1, Pri1 and Pri2 that are likely to promote PIC formation. Indeed, mutations of these residues were found to severely impair the ability of CST to stimulate PP activity *in vitro*. In addition, the corresponding mutations in Stn1 and Ten1 were found to disrupt telomere regulation *in vivo*, resulting in two distinct phenotypes. One class of mutant exhibited progressive telomere lengthening without evidence of telomere DNA damage, whereas the other mutants manifested slow growth and multiple telomere aberrations indicative of telomere deprotection. DNA repair factors including Rad52, Rad24 and Exo1 were shown to modulate the growth and telomere defects in the telomere deprotection mutants. Collectively, our findings support the notion that besides C-strand synthesis, the yeast CST and PP complexes act jointly to inhibit telomerase and protect telomeres, underscoring the multiple functions of a replicative DNA polymerase in telomere regulation.

## Materials and Methods

### *Candida glabrata* strain construction and growth

Standard protocols were employed for the genetic manipulation of *C. glabrata*. All strains used in this study are derived from the BG14 strain background (23) and are listed in Supp. Table 1.

The *exo1Δ, rad52Δ, rad24Δ* and *tertΔ* disruption strains were constructed by integrating an NAT^R^ cassette flanked by ∼600 – 700 bp of 5’ UTR and 3’ UTR derived from the corresponding genes (24). The UTR fragments were generated by PCR amplification and cloned in between the *Sac*II and *Hpa*I sites, as well as between the *Aat*II and *Pvu*II/BsiWI sites of pBV65. The cassettes were then released from the plasmid via *Aat*II and *Hpa*I cleavage and used in transformation. Correct clones were selected on media containing 100 µg/ml Nourseothricin and identified by PCR genotyping. The FLAG_3_-tagged *TEN* gene was constructed by cloning the ORF upstream of the FLAG_3_ tag in between the *Bam*HI and *Not*I sites of pSMT3-FLAG_3_ (25). The tagged gene was then reamplified with appropriate primers and cloned in between the *Xba*I and *Xho*I sites of pCU-PDC1, pCU-HHT2, and pCU-EGD2 plasmids. PCR mutagenesis was used to generate the various Ten1 mutant expression plasmids. The primers used for these constructions are all listed in Supp. Table 2.

For growth analysis of the mutants, liquid cultures of *C. glabrata* strains were first diluted to OD_600_ of 1.0 (corresponding to ∼ 3 x 10^7^ cells/ml) and subjected to serial 6-fold dilutions. Four microliters of the diluted cultures were then spotted onto semi-solid media and grown at 30° C for 1 to 2 days prior to image acquisition.

### Telomere length and structural analyses

Southern analysis of telomere restriction fragments (TRF Southern) was performed using DNA treated with *Eco*RI or *Eco*RI plus *Alu*I as described previously (26). The blots were hybridized to an oligonucleotide containing three copies of the *C. glabrata* C-strand repeat (CgTEL-C3; [GCACCCAGACCCCACA]_3_). For calculation of average TRF lengths, we used the formula mean TRF length = Σ (OD_i_ × L_i_) / Σ (OD_i_) because the majority of TRF length variation stems from differences in subtelomeric lengths (27,28). The in-gel hybridization assays for detecting G-strand ssDNA have been described (29). The dried gels were hybridized to a 24-mer probe (CgTEL-C1.5) corresponding to 1.5 copies of the *C. glabrata* C-strand repeat. C-circle and G-circle assays were performed as before except that deoxynucleotides triphosphates needed for the *C. glabrata* G-strand and C-strand synthesis were used, respectively (30).

### Protein expression and PP activity assays

The *C. glabrata* CST, ST and PP complexes were obtained via recombinant co-expression of the relevant subunits in E. coli and affinity purification. For CST, *E. coli* BL21 (DE3) was co-transformed with pACYC-SUMO-Cdc13-FG (Cdc13 with an N-terminal SUMO and C-terminal FLAG tag) and pCDF-H10ST (Stn1 with an N-terminal His_10_ tag and untagged Ten1). For C*ST, *E. coli* BL21 (DE3) was co-transformed with pDV1BUK-C2-Strep-Cdc13_OB234_ (encoding Strep-Trx-CL7-SUMO-TEV-Strep-Cdc13_OB234_) and pCDF-H10ST. All these plasmids have been described previously (21,26). Mutations in the STN1 and TEN1 ORFs were introduced by PCR mutagenesis using appropriate oligonucleotides (Supp. Table 2). Following induction of the co-transformed *E. coli* clones by IPTG, the CST complex was purified from extracts over a Ni-NTA column, treated with ULP1 to remove the SUMO tag, and then further purified through binding and elution from anti-FLAG Sepharose beads (GenScript Biotech. Corp.). The purification of the C*ST complex was the same except that Strep-Tactin chromatography was used as the second affinity step. For PP, the ORFs of the four subunits of the complex, Pol1, Pol12, Pri1 and Pri2, were amplified from *C. glabrata* lysates using PCR. The polymerase subunits, Pol1 and Pol12, were cloned in MCS1 and 2 of a modified pRSFDuet vector respectively. Similarly, the primase subunits, Pri1 and Pri2, were cloned in MCS1 and 2 of a modified pCDFDuet vector, respectively. For the expression of the complex in bacteria, the *E. coli* BL21 (DE3) strain was co-transformed with pRSF-H10Pol1-Pol12 (Pol1 with an N-terminal His_10_ tag and Pol12 with a C-terminal Strep-tag-II) and pCDF-Pri1-Pri2 (untagged Pri1 and Pri2 with a C-terminal Strep-tag-II). Following extract preparation, the complex was first enriched through a Ni-NTA column, and then further purified over a Strep-Tactin column. The ST complex was expressed using the pCDF-H10Stn1-Ten1FG_3_ plasmid (Stn1 with an N-terminal His_10_ tag and Ten1 with a C-terminal FLAG_3_ tag). This plasmid was made by replacing the Ten1 ORF in pCDF-H10ST with the Ten1-FG_3_ ORF from pCU-PDC1-Ten1-FG_3_ and the complex was purified using Ni-NTA and FLAG affinity chromatography.

The PP activity assays were performed in 20 µl reactions containing 40 mM Tris.HCl, pH 7.6, 30 mM potassium acetate, 8 mM Mg acetate, 5 mM dithiothreitol, 0.05 mM EDTA, 5% glycerol, 0.1 mg/ml bovine serum albumin, ∼5-10 nM PP, CST or ST at specified concentrations, 1 mM ATP, 10 µM dATP (with 0.5 to 1 µCi α-P^32^-dATP), and 300 ng of poly-dT (Midland Inc.) (18). After incubation at 32°C for 60 m, reactions were terminated with 80 µl STOP solution (10 mM Tris.HCl, pH 8.0, 20 mM EDTA) and 100 µl proteinase K solution (10 mM Tris.HCl, pH 8.0, 0.5% SDS, 0.20 mg/ml proteinase K), and incubated at room temperature for 30 m. The products were recovered by ethanol precipitation in the presence of 20 µg GlycoBlue and 30 µg tRNA, resuspended in 10 µl formamide loading dye and analyzed by electrophoresis through a 13% polyacrylamide, 7M urea gel in 1x TBE buffer. Signals were acquired by PhosphorImager scanning (Typhoon FLA 7000) and analyzed using PRISM.

### Structural Predictions

All protein and protein-DNA complex structural predictions were performed using the AlphaFold 3 public server without imposing any post-translational modification and using random seeds (31). We obtained protein sequences from public databases for Pri1 (Uniprot Q6FKQ9), Pri2 (UniProt Q6FLP8), Cdc13 (UniProt Q6FT40), Stn1 (UniProt Q6FM89) and Ten1 (RefSeq XP_065482704.1). The DNA substrate predicted was composed of 2 ¼ repeats of the *C. glabrata* telomeric sequence (5’- [TGTGGGGTCTGGGTGC]_2_TGTG -3’). All predicted alignment error (PAE)(32) plots were created with *af3analysis* (https://github.com/nayimgr/af3analysis). All models were visualized using ChimeraX(33) and low-confidence regions (pLDDT < 30) were removed for easier visualization.

## Results

### Identification of Ten1, Pri1 and Pri2 surface residues involved in CST-mediated Polα stimulation

In the human CST-PP PIC structures, TEN1 is centrally located and makes multiple contacts to PP subunits (Fig. 1a) (22). In particular, the TEN1 L34 (loop between β-strand 3 and 4) wedges between PRIM1 and PRIM2 and makes sequential contacts to a surface loop and a helix on these proteins, respectively (Fig. 1a and 1b). While these surface residues are not well conserved between yeasts and mammals (Supp. Fig. 1), we reasoned that they could have co-evolved to retain interactions in budding yeasts, and proceeded to engineer alanine substitution mutants in *C. glabrata TEN1*, *PRI1* and *PRI2* (designated *ten1^L34A^*, *ten1^L34B^*, *pri1^L34^*, and *pri2^L34^*) aimed at disrupting the hypothesized physical contacts (Supp. Fig. 1). The mutant CST and PP complexes as well as the wild type controls were expressed in *E. coli*, purified through sequential affinity chromatography (Supp. Fig. 2a) and subjected to *in vitro* assays to assess the impacts of mutations on CST-mediated PP stimulation.

**Fig. 1.**
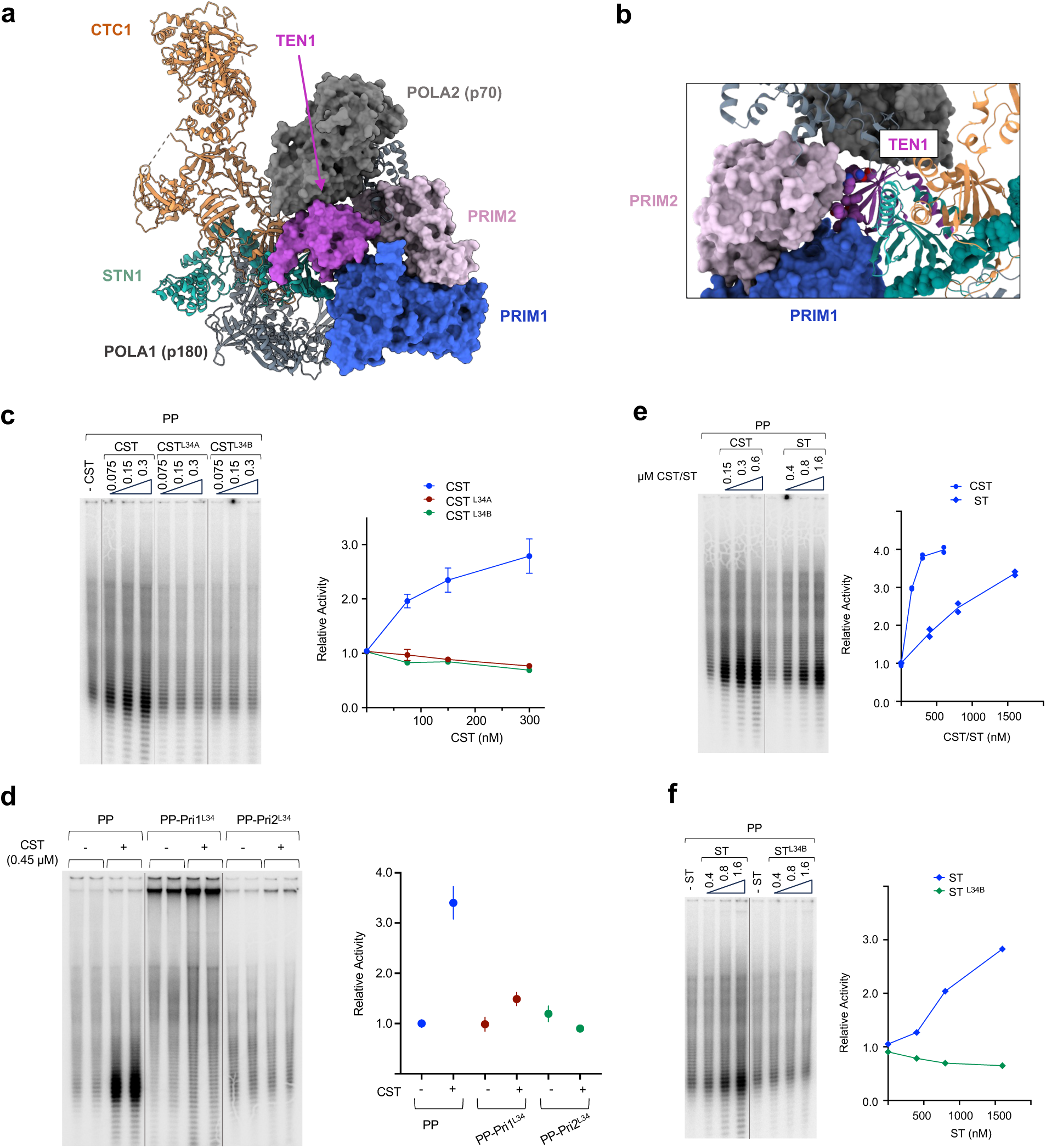
*C. glabrata* Ten1, Pri1, and Pri2 surface loops are required for the stimulation of PP activity by the CST complex *in vitro*. **a.** Structure of the human CST-PP-DNA pre-initiation complex illustrating the central location of TEN1 (magenta) and its contacts to the primase subunits (PRIM1 and PRIM2). **b.** A magnified image of the CST-PP-DNA complex in a different orientation highlighting the roles of the TEN1 L34 loop in mediating interactions with PRIM1 and PRIM2. **c.** The activity of the PP complex in synthesizing poly-rA-poly-dA from the poly-dT template was analyzed in the presence of increasing concentrations of CST, CST^L34A^ and CST^L34B^. The results for assays in one experiment are shown on the left and the quantifications from three independent experiments shown on the right. **d.** The activities of PP, PP-Pri1^L34^, and PP-Pri2^L34^ complex were analyzed in the absence or presence of 0.45 µM CST. The results for duplicate assays in one experiment are shown on the left and the quantifications from two independent experiments shown on the right. **e.** PP activity was assayed in the presence of increasing concentrations of CST and ST. The products from one set of assays are displayed on the left and the quantifications from three independent experiments plotted on the right. **f.** PP activity was assayed in the presence of increasing concentrations of ST and ST^L34B^. The products from one set of assays are displayed on the left and the quantifications from three independent experiments plotted on the right.

Notably, our previous *in vitro* analysis utilized PP purified from *C. glabrata* and CST prepared from a different set of *E.coli* expression plasmids. We therefore first compared the different preparations in their ability to support synthesis of poly-rA-poly-dA products from poly-dT template. Both the native PP complex isolated from *C. glabrata* and the recombinant PP from *E. coli* synthesized labeled products that are ATP- and template-dependent (Supp. Fig. 2b). Both forms of PP were also stimulated by the CST complex to similar extent, indicating that the new expression and purification protocols yielded active protein complexes (Supp. Fig. 2b). In addition, the PP complexes harboring either the Pri1^L34^ or Pri2^L34^ mutant protein (PP- Pri1^L34^ and PP- Pri2^L34^) remained active in the synthesis of poly-rA-poly-dA, indicating that these mutations did not abrogate PP activity (Supp. Fig. 2c).

We next examined the effects of the Ten1, Pri1 and Pri2 mutations on the PP-stimulatory activity of CST. Notably, purified CST complexes harboring the Ten1^L34A^ or Ten1^L34B^ mutant proteins (CST^L34A^ or CST^L34B^) were found to contain the three components at similar ratios as the wild type complex, indicating that the mutations do not affect complex assembly (Supp. Fig. 2a). The mutant complexes also exhibited comparable DNA-binding activity for the *C. glabrata* telomere G-strand as the wild type complex, indicating that they retain as least some biochemical activity (Supp. Fig. 2d). In parallel PP activity assays, we found that whereas wild type CST produced dose-dependent enhancement of PP activity, CST^L34A^ and CST^L34B^ were mostly inactive (Fig. 1c). In addition, even though PP-Pri1^L34^ and PP-Pri2^L34^ retain substantial basal activity in the absence of CST, neither complexes are stimulated in the presence of CST (Fig. 1d). These results suggest that the structural elements implicated in the formation of the human CST-PP PIC complex (22) are also required for the regulatory activity of yeast CST. Hence, despite poor sequence conservation at the Ten1-Pri1-Pri2 interface, the human and yeast CST complexes appear to utilize similar mechanisms to promote PP activity.

### The Stn1-Ten1 subcomplex can stimulate PP at high concentrations

While Stn1 and Ten1 are well conserved in evolution at the level of overall structure, the CTC1/Cdc13 family of proteins exhibit a high degree of evolutionary divergence (7,14,34). Given that PP stimulation is a conserved CST function, we hypothesize that the CTC1/Cdc13 subunit may be dispensable for this activity. To test this idea, we purified a Stn1-Ten1 (ST) subcomplex and compared its stimulatory activity to the full CST complex. Indeed, at high concentrations, ST can stimulate PP to a similar extent as CST (Fig. 1e). In standard assays, ∼7 fold more ST than CST is required to achieve 1/2 maximal stimulation. Notably, like CST, the ST complex harboring Ten1^L34B^ (ST ^L34B^) failed to enhance PP activity, supporting the notion that ST acts via similar mechanisms as the full CST complex (Fig. 1f).

We had previously shown that Stn1 alone is capable of stimulating PP (18). This Stn1-exclusive stimulatory activity, which was observed using Stn1 that carried a His-SUMO tag, is evidently at odds with the Ten1 mutational analysis. One possibility is that the absence of Ten1 may have exposed normally inaccessible portions of Stn1 to engage in fortuitous interaction with PP. Given the consistent effects of Ten1 mutations both *in vitro* and *in vivo* (Fig. 1 and see later), the Stn1-exclusive stimulatory activity is unlikely to accurately reflect the *in vivo* mechanisms of the CST complex.

### AlphaFold modeling supports similarities between the human and *C. glabrata* CST-PP PIC complex

After the above analysis was completed, the AlphaFold 3 (AF3) structure prediction algorithm became available (31). Queries of the AF3 server using *Cg*CST and primase protein subunits along with a 36-mer *C. glabrata* G-strand ssDNA yielded models of the *Cg*CST-primase-DNA complex that resemble the human complex regarding the dispositions of Stn1, Ten1, Pri1, Pri2 and DNA (Supp. Fig. 3a and 3b). In particular, the *Cg*Ten1 L34 loop in the AF3 models is wedged adjacent to the Pri1 and Pri2 interface, just like the human proteins in the cryo-EM structures (Supp. Fig. 3a and 3b). In addition, based on the confidence measure developed for AlphaFold (PAE), the interactions between Ten1 and Pri1/Pri2 are estimated to be comparable to those within individual proteins (Supp. Fig. 3c). Notably, this PAE-based signal for interaction is present in all 5 models generated by AF3, adding confidence to the validity of the structural prediction.

Given the ability of the *Cg*Stn1-Ten1 complex to stimulate PP activity *in vitro*, we also used AF3 to predict the structure of a complex comprising Stn1, Ten1, primase and DNA. Indeed, AF3 generated models of the *Cg*ST-primase-DNA complex that recapitulate the interactions between Stn1, Ten1, Pri1, Pri2 and DNA found in the *Cg*CST-primase-DNA complex (Supp. Fig. 4a). Again in this case, the PAE-based signals for interactions between Ten1 and Pri1-Pri2 are detected in all 5 models generated by AF3, supporting the validity of the structural prediction (Supp. Fig. 4b). Altogether, the results from biochemical analysis and structural modeling support the idea that despite sequence divergence, the yeast and human Ten1, Pri1 and Pri2 orthologs utilize equivalent surface loops to promote PIC formation.

Interestingly, the Ten1-primase interactions in AF3 models of the *Cg*CST-primase-DNA and *Cg*ST-primase-DNA complexes were not predicted by AF3 when the full PP complex (Pri1, Pri2, Pol1 and Pol12) were included in the modeling exercise (not shown). One likely explanation is that because the CST-PP-DNA complex can undergo more conformational changes than CST-primase-DNA, the computational challenges for machine learning-based algorithms are greater (35).

### The *ten1^L34A^* and *ten1^L34B^* mutants induce distinct telomere defects *in vivo*

Having established the importance of the Ten1 L34 loop in stimulating PP activity, we next examined the functions of this loop *in vivo* by over-expressing FLAG_3_-tagged Ten1^L34A^ and Ten1^L34B^ proteins (from a plasmid harboring the strong *PDC1* promoter) (36). This strategy allows us to prioritize mutant proteins that can compete effectively against wild type Ten1 for assembly into the CST complex, but that render the complex defective in telomere regulation. For comparative purposes, we tested three additional mutants with altered residues in other Ten1 surface loops (i.e., Ten1^L12^, Ten1^L23^, and Ten1^L5C^) (Fig. 2a). All the mutant as well as the wild type Ten1 plasmids yielded viable transformants. Western analysis confirmed that the Ten1 alleles encoded by these plasmids were all expressed, but that Ten1^L34B^ and Ten1^L5C^ were present at ∼ 1/5 of the level of wild type, FLAG_3_-tagged Ten1 (Fig. 2b). Given that the *PDC1* promoter is a particularly strong one in *C. glabrata*, it seems likely that even the reduced Ten1^L34B^ and Ten1^L5C^ levels may still be expressed at sufficient levels to compete against endogenous Ten1. Indeed, while most mutants grew at near wild-type growth rate, the Ten1^L34B^-bearing mutant manifested slow growth suggesting defective telomere function (Fig. 2c).

**Fig. 2.**
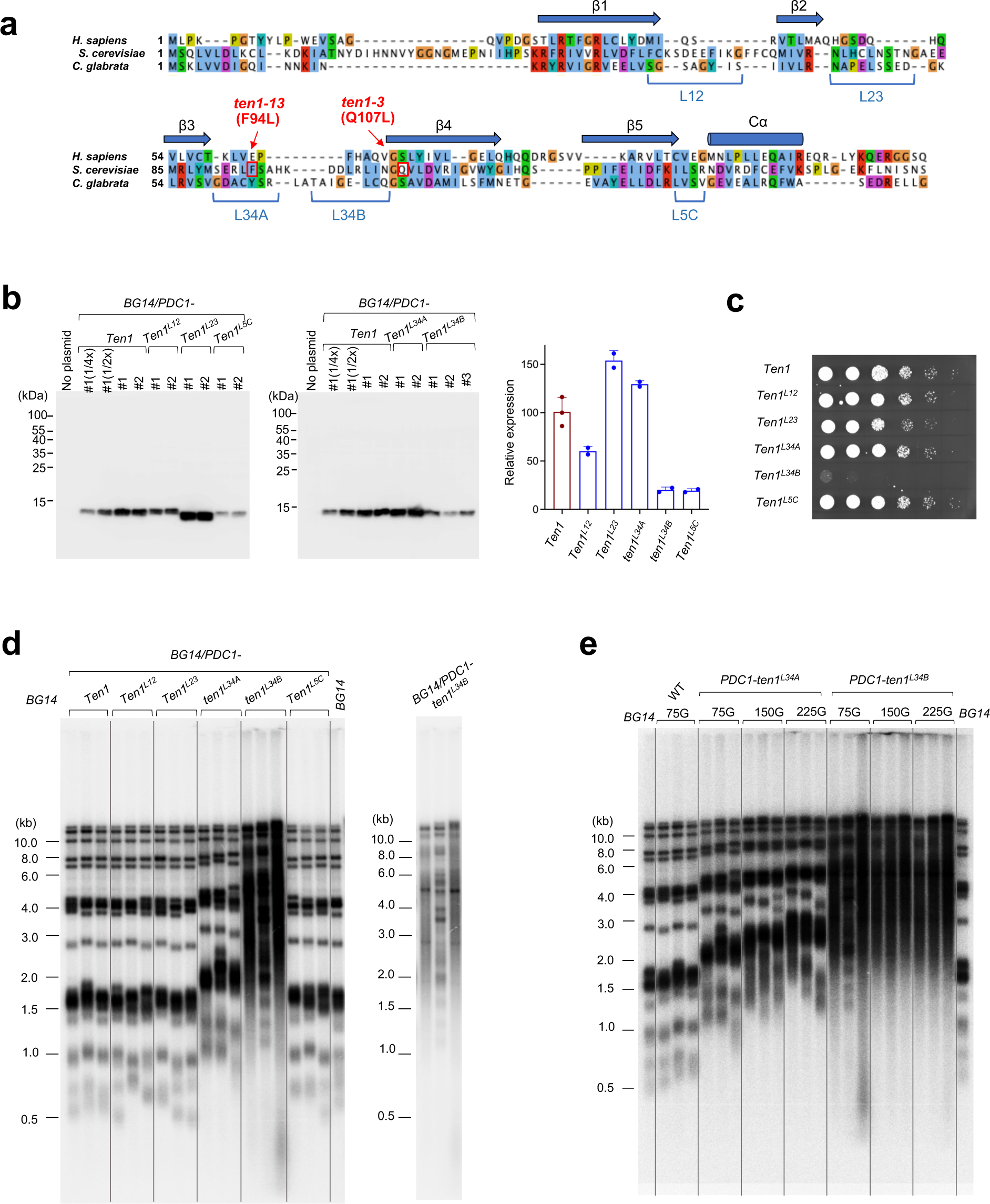
t*e*n1 mutants that impairs the ability of CST to stimulate PP trigger distinct telomere aberrations *in vivo*. **a.** An alignment of the Ten1 orthologs from humans (*H. sapiens*), *S. cerevisiae*, *C. glabrata* illustrating the locations of the five conserved β-strands and C-terminal α-helix. Also indicated are the locations of two temperature-sensitive *Sc*Ten1 mutants (*ten1-13* and *ten1-3*) and the five cluster Alanine *Cg*Ten1 mutants analyzed in this study (L12, L23, L34A, L34B, and L5C). **b.** Western analysis of FG_3_-tagged Ten1 in lysates from clones transformed with the indicated expression plasmids. Duplicated assay results are shown on the left and the quantifications plotted on the right. Note that a dilution series was performed using the one of the clones carrying *PDC1-Ten1* to generate a standard curve for quantitation (2^nd^ to 4^th^ lanes). **c.** Serial dilutions of BG14 clones overexpressing the indicated *TEN1* allele were spotted onto SD-ura semi-solid media and incubated at 30 degree for 1 day. **d.** Genomic DNAs were prepared from three independent transformants carrying the indicated Ten1 allele after ∼75 generations of growth, treated with EcoRI, and subjected to telomere restriction fragment (TRF) Southern. A less saturated display of the *PDC1-ten1^L34B^* samples is shown on the right to better illustrate the increased TRF length heterogeneity. **e.** Genomic DNAs were prepared from cells with either *PDC1-ten1^L34A^* or *PDC1-ten1^L34B^* (three clones each) after the indicated number of generations, treated with EcoRI, and subjected to telomere restriction fragment (TRF) Southern.

Analysis of telomere lengths following ∼75 generations of growth (3 rounds of re-streaking for single colonies) revealed distinct, abnormal telomere patterns in the Ten1^L34A^ and Ten1^L34B^ mutants; the former showed consistent telomere lengthening (∼ 300 bp) for the great majority of their telomere restriction fragments (TRF), whereas the latter showed increased telomere lengths as well as heterogeneity (Fig. 2d). In contrast to Ten1^L34A^ and Ten1^L34B^, overexpression of wild type Ten1 and all other mutants did not result in telomere aberrations. We then monitored telomere dynamics in the Ten1^L34A^ and Ten1^L34B^ cells for another 150 generations and found that the telomere elongation in Ten1^L34A^-expressing cells is progressive, at a rate of ∼ 4 bp per generation, while the telomere abnormalities triggered by Ten1^L34B^ occurred early and did not exacerbate over time (Fig. 2e). These initial studies were performed using DNAs treated with *Eco*RI alone, which resulted in multiple TRF clusters owing to the different lengths of the subtelomeric DNA from different chromosomes. To simplify the analysis and validate the initial finding, we repeated the TRF measurement using DNAs treated with both *Eco*RI and *Alu*I, which resulted in a single predominant TRF cluster (∼1.2 kb average lengths in the wild type strain) (Supp. Fig. 5a). Analysis using this protocol again revealed progressive telomere elongation in Ten1^L34A^ cells and stably elongated, heterogeneous telomeres in Ten1^L34B^ cells (Supp. Fig. 5a and 5b). Interestingly, after prolonged passage, the Ten1^L34A^ telomere TRFs appear to stabilize at ∼ 3.0 kb (Supp. Fig. 5c), whereas Ten1^L34B^ TRFs showed a mild decline (Supp. Fig. 5b). We conclude that even though both mutations impaired CST-PP interaction, they result in different abnormalities *in vivo*.

### The *ten1^L34A^* mutant is defective in telomerase inhibition *in vivo*

The progressive telomere elongation triggered by Ten1^L34A^ suggests aberrant telomerase regulation, which has been reported for a number of CST mutants (37-39). To confirm the telomerase-dependence of telomere elongation induced by Ten1^L34A^, we first generated several independent *tertΔ* clones (missing the catalytic component of telomerase) and characterized their telomere dynamics. Consistent with previous findings (40), we found that *tertΔ* cells manifest progressive telomere shortening, followed by senescence and emergence of survivors after ∼ 150 generations of growth (Fig. 3a). We then overexpressed Ten1 and Ten1^L34A^ in early generations of *tertΔ* clones and analyzed telomeres after ∼35 generations of growth. Notably, while Ten1^L34A^ expression induced longer telomeres than Ten1 in the parental strain (BG14), it failed to do so in *tertΔ* cells, indicating that the abnormal telomere lengthening requires telomerase (Fig. 3b).

**Figure 3.**
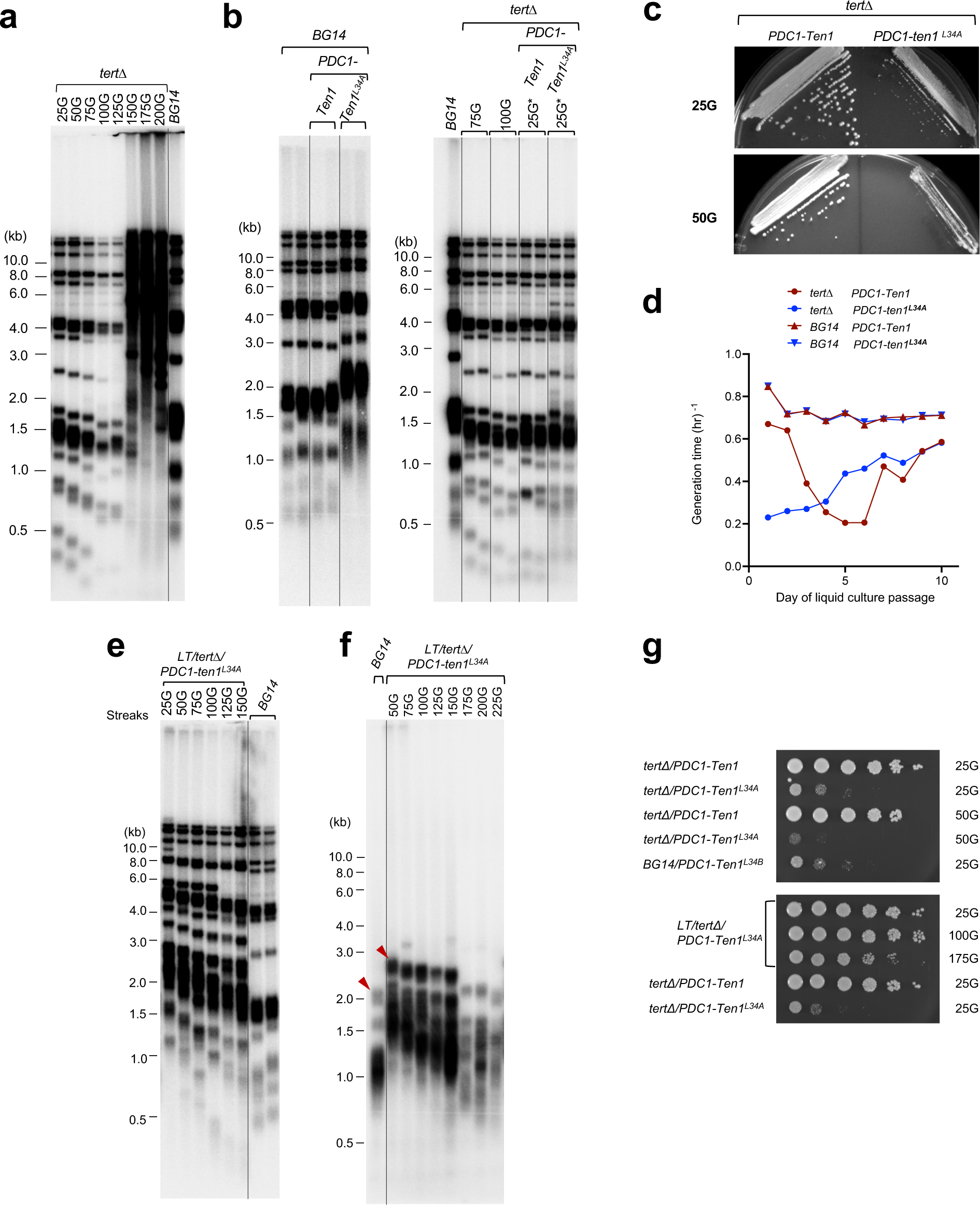
Overexpression of the Ten1^L34A^ mutant protein results in telomerase-dependent telomere elongation and causes accelerated senescence in cells with short telomeres. **a.** Genomic DNAs were prepared from a freshly constructed *tertΔ* clone (from BG14) after the designated number of generations, cleaved with *Eco*RI, and then subjected to TRF Southern analysis. **b.** The BG14 and *tertΔ* strains were transformed with plasmids expressing *PDC1-Ten1* or *PDC1-ten1^L34A^*. Genomic DNAs were prepared after the indicated number of generations, cleaved with *Eco*RI, and subjected to TRF Southern analysis. Note that the *tertΔ*/*PDC1-Ten1* and *tertΔ*/*PDC1-ten1^L34A^* DNAs were prepared from cells that had been grown for 25 generations after the introduction of the Ten1 expression plasmids (25*), but ∼ 75 generations after the disruption of *TERT*. **c.** The *tertΔ*/*PDC1-Ten1* and *tertΔ*/*PDC1-ten1^L34A^* clones were grown for ∼25 and 50 generations after the introduction of the Ten1 expression plasmids and then streaked on semi-solid media to monitor senescence. **d.** The indicated strains were continuously passaged by in liquid culture, and the generation time for each day estimated from the population doubling for the overnight cultures (see Materials and Methods). **e.** The *BG14*/*PDC1-ten1^L34A^* strain was grown for 225 generations to create a clone with long telomeres (*LT/PDC1-ten1^L34A^*), and then transformed with the TERT disruption cassette. The resulting strain (*LT/tertΔ/PDC1-ten1^L34A^*) was passaged for the indicated number of generation, followed by genomic DNA preparations. The DNAs were cleaved with *Eco*RI and subjected to TRF Southern. **f.** Same as E except that genomic DNAs were treated with *Eco*RI and *Alu*I prior to TRF Southern. **g.** Serial dilutions of the indicated strains were spotted onto SD-ura semi-solid media and incubated at 30 degree. Images of the plates were collected after 2 days of incubation. The numbers of cell divisions the strains have undergone following *TERT* disruption and the introduction of the Ten1 plasmids are indicated on the right.

Interestingly, the *tertΔ/PDC1-ten1^L34A^* cells exhibited rapid onset of senescence (at ∼25G post construction) despite having comparable telomere lengths as *tertΔ/PDC1-Ten1* cells (Fig. 3b and 3c). Continuous liquid cultures of *tertΔ/PDC1-ten1^L34A^* cells resulted in the emergence of faster growing survivors, just like *tertΔ/PDC1-Ten1* cells (Fig. 3d). The accelerated senescence and survivor formation for *tertΔ/PDC1-ten1^L34A^* cells suggests that the Ten1^L34A^ may induce partial telomere deprotection when telomeres are short. To examine this issue further, we asked whether expression of *Ten1^L34A^* in *tertΔ* strain with long telomeres can also trigger rapid senescence. This was accomplished by disrupting the *TERT* gene in a *C. glabrata* clone that had harbored the Ten1^L34A^ expression plasmid for 225G. This LT/*tertΔ*/*PDC1-Ten1^L34A^* (long telomere with *tertΔ* and *PDC1-Ten1^L34A^*) strain possessed telomeres that are ∼800 bp longer than the parental strain (BG14) shortly after *TERT* deletion (Fig. 3e, compare the TRFs marked by arrowheads). *LT/ PDC1-Ten1^L34A^/tertΔ* showed a similar telomere attrition rate as *tertΔ,* confirming that the telomere elongation induced by Ten1^L34A^ is telomerase-dependent (Fig. 3e and 3f). Notably, in contrast to *tertΔ/PDC1-ten^L34A^*, *LT/ tertΔ/PDC1-ten^L34A^* can be passaged for more than 150G without growth defects (Fig. 3g). Hence, the rapid senescence exhibited by *tertΔ/PDC1-ten^L34A^* appears to be restricted to cells with short telomeres.

### The *ten1^L34B^* mutant induces telomere deprotection, leading to DDR and aberrant repair

In contrast to *ten1^L34A^*, the slow growth and telomere length heterogeneity exhibited by *ten1^L34B^*-O/E cells suggest loss of telomere protection. Consistent with this interpretation, we found telomeres in these cells to be more heterogeneous than those in post-senescent survivors, which utilize a recombination-based telomere maintenance pathway (Supp. Fig. 6a). Specifically, while discreet TRF clusters can be discerned within the broad telomere smears for survivors, no such clusters were observed for *ten1^L34B^* cells. The *ten1^L34B^* mutant also manifested elevated G-strand ssDNA and C-circles, which are again hallmarks of telomere deprotection (Supp. Fig. 6b and 6c).

We next sought to assess the roles of specific DDR and DNA repair factors, including Rad52, Rad24 and Exo1, in mediating the defects of the *ten1^L34B^* mutant. These genes were chosen for their strong connections to dysfunctional telomeres in budding yeast (41-43). To alleviate the challenges posed by the toxicity of Ten1^L34B^, we constructed Ten1^L34B^ expression plasmids carrying the *HHT2* or *EGD2* promoter, which were reported to be weaker promoters than *PDC1* (36). Based on Western analysis, these promoters expressed Ten1 at ∼ 1/5 and 1/10 of the level produced from *PDC1* (Supp. Fig. 7a). As expected, Ten1^L34B^ expressed from plasmids with different promoters suppressed growth in a dose-dependent manner, further confirming its toxic effect (Supp. Fig. 7b). We then constructed isogeneic *rad52Δ*, *rad24Δ*, and *exo1Δ* mutants and examined the consequences of expressing wild type and mutant Ten1 with respect to growth and telomere aberrations. For these experiments, Ten1 and Ten1^L34A^ were expressed from the *PDC1* promoter, whereas Ten1^L34B^ was expressed using either the *HHT2* or the *EGD2* promoter.

We first assessed the effects of *rad52Δ*, *rad24Δ*, and *exo1Δ* on the growth of Ten1-expressing cells on semi-solid media. While none of the deletions caused changes in the growth of cells harboring *PDC1-Ten1*, deletion of *EXO1* substantially improved the growth of cells with *HHT2-ten1^L34B^* and *EGD2-ten1^L34B^,* underscoring the role of this nuclease in mediating the toxic effects of Ten1^L34B^ (Fig. 4a and Supp. Fig. 7c). We then analyzed the effects of *rad52Δ*, *rad24Δ*, and *exo1Δ* on the telomere aberrations triggered by Ten1^L34B^, including telomere lengthening/heterogeneity, ssDNA on the G-strand, and C-circle accumulation (Fig. 4b – 4d). Notably, each deletion had disparate impacts on distinct telomere abnormalities, suggesting that all three genes are partly responsible for acting at deprotected telomeres. In particular, *rad24Δ* reduced telomere elongation without eliminating the increased telomere heterogeneity, while *rad52Δ* and *exo1Δ* had little effects on telomere length distributions (Fig. 4b). With regard to G-strand ssDNA, *rad52Δ* triggered a significant increase in this structure, whereas *rad24Δ* and *exo1Δ* caused mild reductions, which are not statistically significant (Fig. 4c). All three deletions, however, decreased the levels of C-circles, which are believed to be markers of telomere DNA damage and telomere recombination (Fig. 4d) (44,45). Together, these observations support the notion that the telomere aberrations in Ten1^L34B^ cells are produced by the activities of multiple DDR and DNA repair factors.

**Figure 4.**
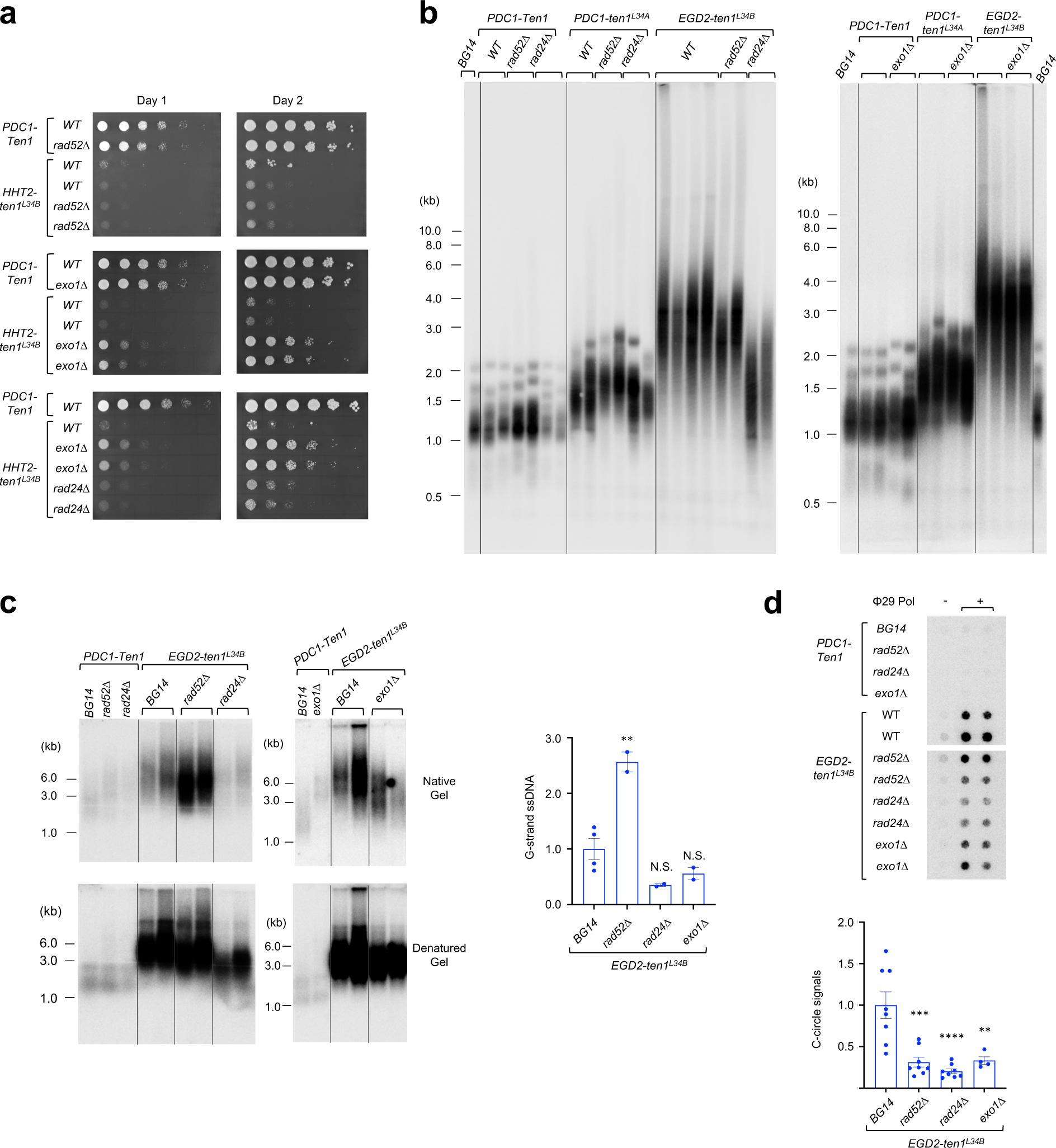
The growth and telomere aberrations of Ten1^L34B^-overexpressing cells are modulated by deletions of DDR genes including *RAD52*, *RAD24* and *EXO1*. **a.** Serial dilutions of the indicated strains were spotted onto SD-ura semi-solid media and incubated at 30 degree. Images of the plates were collected after 1 and 2 days of incubation. **b.** Independent clones of the indicated strains were passaged for ∼75 generations following the introduction of the Ten1 expression plasmids. Genomic DNAs were prepared, digested with *Eco*RI and *Alu*I, and then subjected to TRF Southern analysis. **c.** Same as B except that the genomic DNAs treated with *Eco*RI and *Alu*I were subjected to in-gel hybridization analysis for G-strand ssDNA. The gels were first hybridized to the probe without alkaline denaturation. After acquisition of the ssDNA signals, the samples in the gels were denatured and rehybridized to the same probe. Representative images from the first and second round of hybridizations are shown on the left and the quantification plotted on the right. P-values are designated by asterisks (**, P < 0.05). **d.** Same as B except that genomic DNAs treated with *Eco*RI were subjected to C-circle analysis. Representative assays are shown on top, and the quantification plotted on the bottom. P-values are designated by asterisks (**, P < 0.05; ***, P < 0.01; ****, P < 0.001).

### The DNA-binding residues of Stn1 are also required for CST-mediated PP stimulation and for telomere protection

Our analysis of Ten1 mutants suggests that formation of the yeast CST-PP-DNA PIC complex is critical for both telomerase inhibition and telomere protection. To further assess this hypothesis, we examined another interaction within this complex, that between Stn1 and DNA. In the cryo-EM structure and AlphaFold model, Stn1 not only binds Ten1 (as predicted from numerous prior studies (46,47)), but also contacts telomere DNA before its entry into the primase complex (Supp. Fig. 8a and 8b) (22). The Stn1-DNA interaction may thus cooperate with other protein-protein contacts to promote PIC formation. In support of this idea, we found that omitting the DNA substrate completely abolished the PAE signals between Ten1 and the Pri1-Pri2 proteins in AF3 models of the *Cg*CST-primase or the *Cg*ST-primase complexes (*cf* Supp. Fig. 8c and Supp. Fig. 3c; *cf* Supp. Fig. 8d and Supp. Fig. 4b). To interrogate the importance of Stn1-DNA interaction experimentally, we designed two Stn1 mutants (Stn1^KK^: K69E, K74E; Stn1^DBM^: K69E, K74E, F127E) that, based on previous studies, are predicted to be DNA-binding deficient (21,48). The ST complexes harboring these Stn1 mutations were purified and tested for their ability to stimulate PP (Fig. 5a and 5b). Consistent with the importance of Stn1 DNA-binding, neither S^KK^T nor S^DBM^T stimulated PP activity even at the highest concentrations (Fig. 5b). We also tested Stn1^KK^ as part of the CST complex. For this analysis, we developed an alternative expression strategy for CST to improve yield and homogeneity. Specifically, because full length Cdc13 is prone to proteolysis between OB1 and OB2, we expressed Cdc13^OB234^ (with an N-terminal Strep-CL7-SUMO tag to facilitate purification) along with His-Stn1 and Ten1 and purified the complex by affinity chromatography (Supp. Fig. 9a). We verified that this complex with truncated Cdc13 but otherwise wild type sequences (named C*ST) produced dose-dependent stimulation of PP activity (Supp. Fig. 9b). By contrast, the C*S^KK^T complex exhibited essentially no stimulatory activity (Supp Fig. 9b). We conclude that the DNA-binding activity of Stn1 is also important for PP regulation.

**Figure 5.**
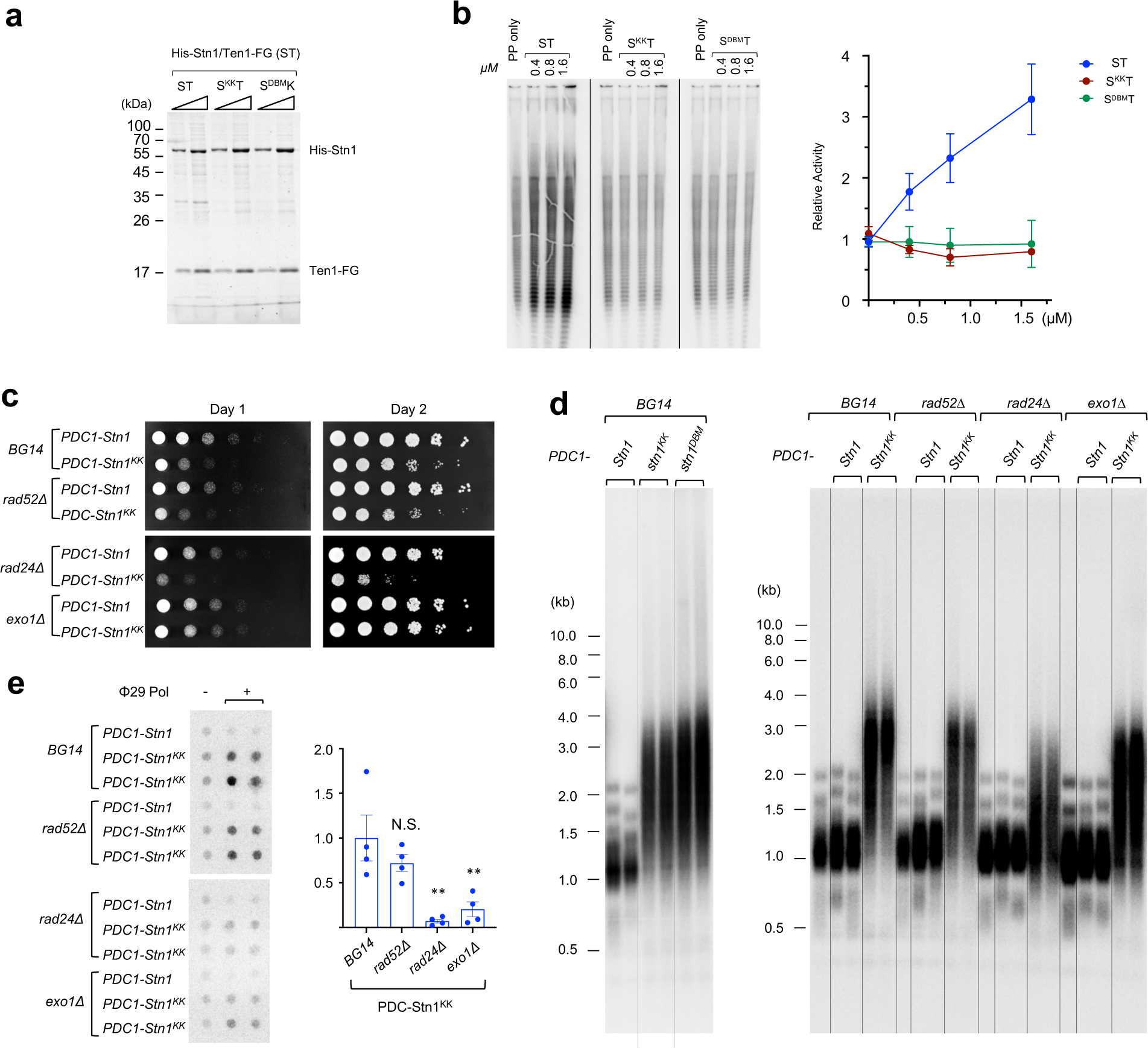
Disrupting Stn1-DNA interaction impaired the stimulatory activity of CST on PP, and caused growth defects as well as telomere aberrations. **a.** Purified ST, S^KK^T, and S^DBM^T complexes were subjected to SDS-PAGE and Coomassie staining. **b.** PP activity was assayed in the presence of increasing concentrations of ST, S^KK^T, and S^DBM^T. The products from one set of assays are displayed on the left and the quantifications from three independent experiments plotted on the right. **c.** Serial dilutions of the indicated strains were spotted onto SD-ura semi-solid media and incubated at 30 degree. Images of the plates were collected after 1 and 2 days of incubation. **d.** Independent clones of the indicated strains were passaged for ∼75 generations following the introduction of the Stn1 expression plasmids. Genomic DNAs were prepared, digested with *Eco*RI and *Alu*I, and then subjected to TRF Southern analysis. **e.** Same as D except that genomic DNAs were treated with EcoRI and subjected to C-circle analysis. Representative assays are shown on the left and the quantification displayed on the right. P-values are designated by asterisks (**, P < 0.05).

Next, we examined the *in vivo* consequences of over-expressing the Stn1^KK^ mutant protein. Like Ten1^L34B^, the expression of Stn1^KK^ from the *PDC1* promoter resulted in slow growth as well as elongated and heterogenous telomeres, consistent with telomere DDR and abnormal repair (Fig. 5c and 5d). Notably, the magnitudes of the growth and telomere defects for Stn1^KK^ are reduced in comparison to those for Ten1^L34B^, suggesting a milder defect (compare Fig. 5c with 4a; Fig. 5d with 4b). To gain more insights on the nature of telomere deprotection, we again assessed the effects of *rad52Δ*, *rad24Δ*, and *exo1Δ* on the growth and telomere aberrations of Stn1^KK^-expressing cells. This analysis revealed marked similarities in the abilities of these deletions to alter phenotypes of the Stn1^KK^ and Ten1^L34B^-containing cells. For example, among the deletions, only *exo1Δ* substantially improved growth, and only *rad24Δ* reduced the overall lengths of telomeres in Stn1^KK^ cells, just as in the case of Ten1^L34B^ cells (Fig. 5c and 5d). In addition, like Ten1^L34B^, all three deletions decreased the levels of C-circles in the Stn1^KK^ cells, albeit that the reduction in *rad52Δ* is not statistically significant (Fig. 5e). Together, these observations support the notion that Stn1^KK^ induces telomere deprotection through similar mechanisms as Ten1^L34B^, i.e., by disrupting CST-PP assembly.

## Discussion

In the current study, we identified multiple structural features in *C. glabrata* Stn1, Ten1, Pri1 and Pri2 that are required for PP activity stimulation by CST *in vitro*, as well as for telomerase inhibition and telomere protection *in vivo*. This discovery suggests that PP plays an integral role in the two major functions previously ascribed to the yeast CST complex (Fig. 6). Our observations have notable implications for the mechanisms, functions and evolution of the CST-PP complex, which are discussed below.

**Figure 6.**
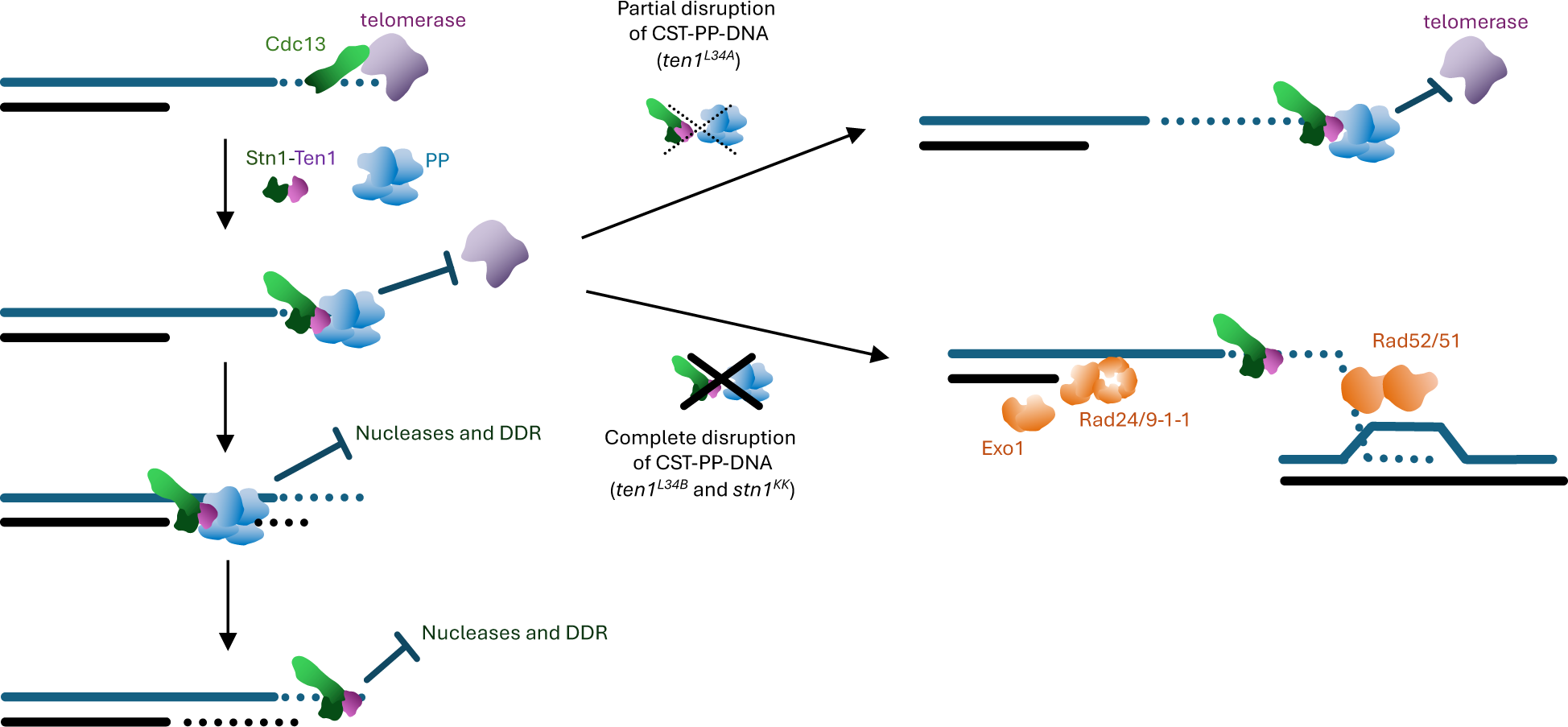
Model for the mechanisms of CST-PP in telomerase termination and telomere protection. Summary model for the switch from G- to C-strand synthesis (left) and the consequences of disrupting CST-PP interactions during this process (right). G-strand extension entails the interactions between Cdc13 and the telomerase complex. Subsequently, the loading of the ST and PP complexes on telomeric DNA results in the termination of G-strand synthesis and the formation of the pre-initiation complex (PIC) for C-strand synthesis. The formation of PIC and the subsequent fill-in synthesis are critical for telomere protection during and following the late S/G2 phase of the cell cycles. Partial disruption of CST-PP-DNA interactions results in delayed PIC formation, allowing telomerase to extend G-strand beyond the normal lengths. Complete disruption of CST-PP-DNA interactions results in the failure to PP to bind telomeric DNA and to mediate C-strand synthesis. Both the absence of PP and the persistent ssDNA on the G-strand renders telomeres susceptible to DDR and aberrant DNA repair.

### Conservation of CST-PP mechanisms

As discussed earlier, the involvement of the Ten1 L34 loop and the Stn1 DNA-binding surface in CST-PP-DNA interaction is first suggested by the disposition of these structures in the human CST-PP pre-initiation complex (PIC). That the equivalent structures in yeast CST are found to be crucial for PP stimulation (despite sequence divergence) suggests that the mechanisms by which CST regulates PP activity is quite well conserved with regard to the formation of PIC. CST likely enhances the initiation of RNA priming by favoring an open conformation of PP, in which the primase subunits are poised to capture the DNA template through Ten1-Pri1-Pri2, Stn1-Ten1, and Stn1-DNA interactions (22,49). Less clear are how CST regulates other steps of the PP reaction and the extent of conservation. The human CST-PP complex at the “recruitment” stage revealed critical contacts mediated by domains of CTC1 that are not conserved in Cdc13, suggesting that the mechanisms of PP recruitment are likely to be different between metazoans and budding yeasts (50). According to a cryo-EM structure of the *Tetrahymena* CST-PP-DNA complex that is proposed to represent the conformation at the DNA polymerization stage, the CST subunits continue to engage in protein-protein interactions with PP (51). Importantly, some of the physical contacts revealed in this structure (e.g., interactions between Pol1 and the Cdc13/Stn1 orthologs) are not observed in the human CST-PP-DNA PIC, suggesting alternative modes of regulation. It is also important to point out that not all physical contacts revealed by cryo-EM structures are functionally relevant. For example, the human TEN1 L23 loop is in close proximity to POLA2 in the PIC structure, yet the corresponding loop in *C. glabrata* Ten1 can be mutated without triggering any apparent defect (Fig. 2c and 2d). Future structural studies coupled with functional analyses in a variety of systems will be needed to provide a more complete understanding of how CST regulates different steps of the PP reaction.

### Termination of telomerase-mediated G-strand synthesis requires the assembly of the CST-PP complex

Human CST is known to inhibit telomerase activity *in vitro* and multiple yeast Cdc13, Stn1, and Ten1 mutations have been shown to induce telomerase-dependent telomere elongation (52,53). However, the molecular mechanisms that underlie CST-mediated telomerase inhibition remain unclear. The biochemical and cellular defects of the *C. glabrata ten1^L34A^* mutant suggests that PP plays an integral role in this pathway, i.e., telomerase inhibition requires the assembly of the full CST-PP complex. First, the *ten1^L34A^* mutant exhibits progressive, telomerase-dependent telomere elongation, consistent with a defect in telomerase termination. Second, the Ten1^L34A^ mutant protein specifically renders the CST complex defective in PP activity stimulation, but has little impact on CST assembly or its DNA-binding activity. Indeed, the interaction with PP represents the only physical contacts for the Ten1 L34 loop observed in all available structures of the CST complex (22,49-51,54). Taken together, these lines of evidence argue that the defect in telomerase termination exhibited by the *ten1^L34A^* mutant is due to disruption of CST-PP.

The accumulated data in budding yeasts suggest that the switch from G- to C-strand synthesis entails the switch from Cdc13-Est1 to Cdc13-Stn1 interaction, with the former supporting telomerase-mediated G-strand elongation and the latter supporting PP-mediated C-strand synthesis (55,56) (Fig. 6). The failure of the Ten1^L34A^ mutant to efficiently terminate G-strand elongation argues that the interaction between Cdc13 and Stn1 by itself is insufficient to remove telomerase, but that the assembly of the full CST-PP complex is required. The coupling of telomerase termination to the assembly of the CST-PP complex ensures that C-strand synthesis can commence immediately following the removal of telomerase. This may in turn maximize the efficiency of converting the newly synthesized G-strand to double-stranded telomeres.

Several observations suggest that the CST-PP mediated telomerase inhibition is conserved in yeasts and mammals. First, two *S. cerevisiae ten1* mutants that were previously shown to induce telomere elongation (*ten1-3* and *ten1-13*) harbor mutations in or near the L34 loop (Fig. 2a) (39). These mutations probably cause defects also by disrupting CST-PP interaction. Second, many *ts* mutants of *S. cerevisiae* and *S. pombe* PP subunits manifest telomere elongation at permissive or semi-permissive temperatures (15), which could be explained by the failure to form adequate levels of the CST-PP complex. Moreover, a human CTC1 mutation that disrupts Polα interaction as well as a mouse *POLA1 ts* mutant have been shown to induce telomerase-dependent telomere elongation, suggesting conservation of this pathway in mammalian cells (57,58). Notably however, a study in colon cancer cells revealed a lack of requirement for TEN1 in terminating telomerase activity (59). Further investigations will be required to establish the extent of mechanistic conservation.

### Yeast CST engages PP to protect telomeres against degradation and recombination

Yeast CST has long been known to mediate an essential protective function at telomeres (60). All three subunits of the complex are required for viability, and many mutant alleles cause dramatic telomere length alterations as well as accumulation of ssDNA, which arises through DDR-induced telomere degradation and recombination (39,42,61-63). The protective function of CST has been ascribed to direct binding of this complex to G-tails to prevent DNA damage recognition and aberrant repair. However, two key findings from our analysis of the *ten1^L34B^* and *stn1^KK^* mutants indicate that CST does not function alone, but rather enlists PP to provide full protection. First, we showed that both *ten1^L34B^* and *stn1^KK^* abolished CST-PP interaction without affecting CST assembly or its DNA-binding activity. Second, we showed that the telomere phenotypes induced by *ten1^L34B^* and *stn1^KK^* resemble previously described CST mutants and are suppressed by the same DNA repair gene deletions such as *exo1Δ* (42,61), implying that they have similar mechanistic basis.

Even though a role for PP in telomere protection has not received much attention, this notion is supported by many previous observations. For example, both yeast and mouse PP mutants can accumulate ssDNA independent of telomerase, consistent with defects in telomere protection or lagging strand synthesis (15,57). It has also been shown that the protective function of Cdc13 is especially critical during replication and dispensable in other phases of the cell cycle (41); this could be rationalized by Cdc13’s interaction with PP in S phase. Indeed, protection is likely mediated through a combination of two mechanisms: 1) masking of ssDNA and DNA ends by the CST-PP assembly; and 2) conversion of G-strand ssDNA to ds telomeres. Defects in either mechanism can lead to the accumulation of structures that trigger DDR and DNA repair. It should be noted that CST-PP acts not only at telomeres following extension by telomerase, but also following telomere replication on the lagging strands (3). Failure to mask the ssDNA or convert ssDNA to dsDNA on the lagging strands can induce DNA damage just as they do at G-strand synthesized by telomerase.

It is unclear why *ten1^L34B^* and *stn1^KK^* trigger telomere deprotection while *ten1^L34A^* causes telomere elongation, even though all three mutants disrupt CST-PP interactions *in vitro*. One distinction between the *in vitro* and *in vivo* analyses is that the *C. glabrata* CST and PP complexes used in the biochemical assays were purified from bacteria and may lack important post-translational modifications. If these modifications facilitate CST-PP interaction, then the *in vitro* analysis will not accurately reflect the *in vivo* activities of these complexes. We suggest that *in vivo*, the *ten1^L34A^* mutant may retain partial ability to mediate CST-PP interaction and support C-strand synthesis. Perhaps the weaker interaction of CST^L34A^ and PP results in delayed complex formation, allowing telomerase to synthesize longer G-strands (Fig. 6). The notion that *ten1^L34A^* and *ten1^L34B^*/*stn1^KK^* represent mutants of different severities is consistent with the phenotypes of *ten1 ts* mutants (38,39). These mutants often exhibit progressive telomere elongation (*ten1^L34A^*-like) at permissive temperatures, but manifest telomere deprotection (*ten1^L34B^* and *stn1^KK^*-like) at non-permissive temperatures, suggesting that partial and complete disruption of the same activities can give rise to disparate phenotypes (Fig. 6).

### Implications for other CST-PP functions and evolution of the CST complex

As noted in the introduction, PP plays critical roles at replication origins and in the synthesis of Okazaki fragments on the lagging strands. Current evidence suggests that CST does not participate in these processes. Instead, PP is directed to the replisome through interactions with CTF4 and the CMG complex (10,11). Notably, these interactions do not appear to involve the Pri1/Pri2 surface structures that engage with Ten1 in the CST-PP-DNA PIC structure, indicating that the replisome employs very different mechanisms to regulate PP. Instead of directly stimulating RNA priming, the replisome may simply recruit PP to the relevant sites on the template to enable primer synthesis.

Besides telomeres, PP has been implicated in two other CST related functions: double-strand break processing and resistance to replication stress. At double strand breaks, CST-PP is proposed to counteract resection by carrying out fill-in synthesis under the control of 53BP1, RIF1 and shieldin (64,65). Under conditions of replication stress, CST is reported to enhance dormant origin firing during recovery, possibly by stimulating PP activity (66). However, additional mechanisms that invokes interactions between CST and RAD51/MRE11 have also been described (67-69). Importantly, it is not known whether CST-PP employs the same mechanisms at DSBs and sites of replication stress as it does at telomeres. This issue may be addressed by testing the effects of the Ten1, Pri1 and Pri2 mutants we identified on their functions in DSB-related pathways.

As a critical component of the telomere maintenance and protection machinery, the CST complex appears to be extraordinarily malleable, especially regarding the CTC1/Cdc13 subunit. CTC1 is likely to represent the more ancestral form of this protein family given its widespread distribution in metazoans and basal branches of fungi (7,14). In contrast, Cdc13 is restricted to the Saccharomycotina subphylum of budding yeasts, and itself manifests a high degree of structural plasticity (34). Notably, in many fungal branches, including Basidiomycetes and fission yeasts, a plausible CTC1- or Cdc13-like protein has not been identified despite extensive efforts. While it is possible that distantly related proteins in these organisms may have escaped detection, our findings suggest a plausible explanation for how the CTC1/Cdc13 protein may become dispensable. Most notably, we show that high concentrations of ST can stimulate PP activity to the same degree as the CST complex. This *in vitro* observation echoes previous descriptions of a “Cdc13-independent” pathway of telomere protection. In particular, the *S. cerevisiae* Cdc13 was found to be dispensable in cells that over-express Stn1 and Ten1, and the viability of these *cdc13* null cells requires normal Pol12 activity (70). Other viable Cdc13Δ mutants that do not require Stn1/Ten1 overexpression have also been described (62,71). Collectively, these *in vitro* and *in vivo* data suggest that the essential function of fungal Ctc1/Cdc13 may be to facilitate the formation of the ST-PP-DNA assembly by providing additional protein-DNA and protein-protein interactions. Accordingly, it is possible that organisms lacking Ctc1/Cdc13 may have adopted alternative means of accomplishing this task, e.g., by expressing high levels of Stn1-Ten1, by enhancing the affinity of Stn1-Ten1 for DNA, or by using other telomere proteins to recruit ST-PP. Indeed, a recent study of *S. pombe* implicates Pot1 and Tpz1 as key recruiters of ST and PP in telomere replication (72). Further analysis of the ST complexes and their interactions with PP in organisms without Ctc1/Cdc13 should be informative.

## Data Availability

The data generated and/or analyzed during this study are included within the paper and its Supplementary Information or are available from the corresponding author upon reasonable request.

## Acknowledgments

We thank Cecile Fairhead for *C. glabrata* strains and Sophie Huang for performing several spotting assays. We acknowledge Oscar Llorca for advice and supervision of Nayim González-Rodríguez and Javier Coloma in their work for this manuscript. We thank Titia de Lange for discussions and Bill Holloman and Michael O’Donnell for comments on the manuscript. The authors declare no competing interests.

