## Supplementary Figures and Tables for "The yeast CST and Polα/primase complexes act in concert to ensure proper telomere maintenance and protection"

**Supplementary Figures 1 – 9**

**Supplementary Tables 1 – 2**

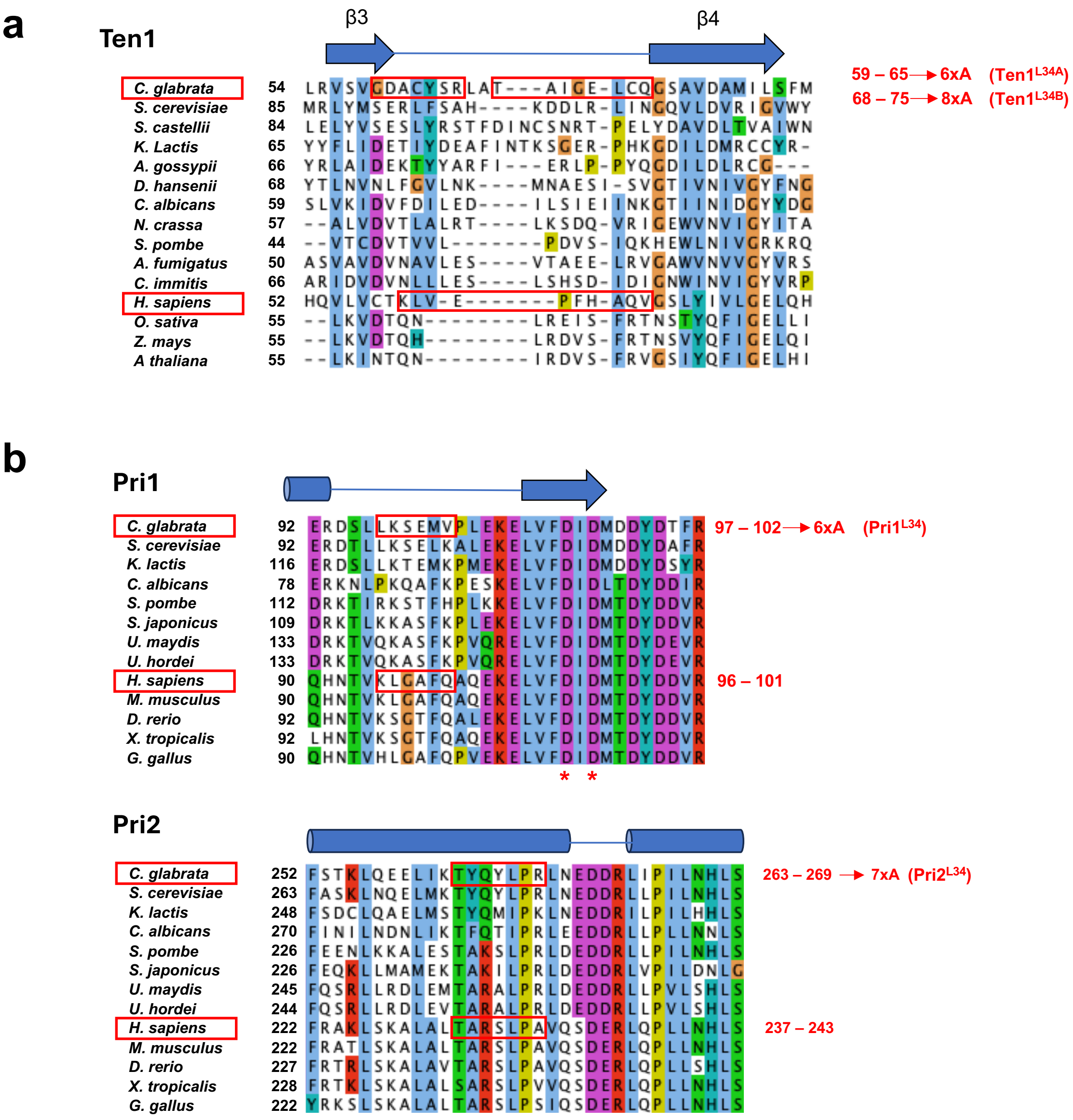

**Supp. Fig. 1. Multiple sequence alignment of the surface loop or helix engaged in the Ten1-Pri1-Pri2 three-way interactions in the CST-PP-DNA PIC complex**

**a.** and **b.** Multiple sequence alignments of the regions of Ten1, Pri1 and Pri2 that engage in protein-protein interactions in the CST-PP-DNA complex. The secondary structures of the aligned regions are illustrated at the top of each alignment. The sequence segments engaged in the three-way interaction are designated by red rectangles with the amino acid residue numbers shown on the right. The cluster Alanine mutations introduced into the *C. glabrata* Ten1, Pri1, and Pri2 proteins are also designated by red rectangles with the amino acid residue numbers shown on the right.

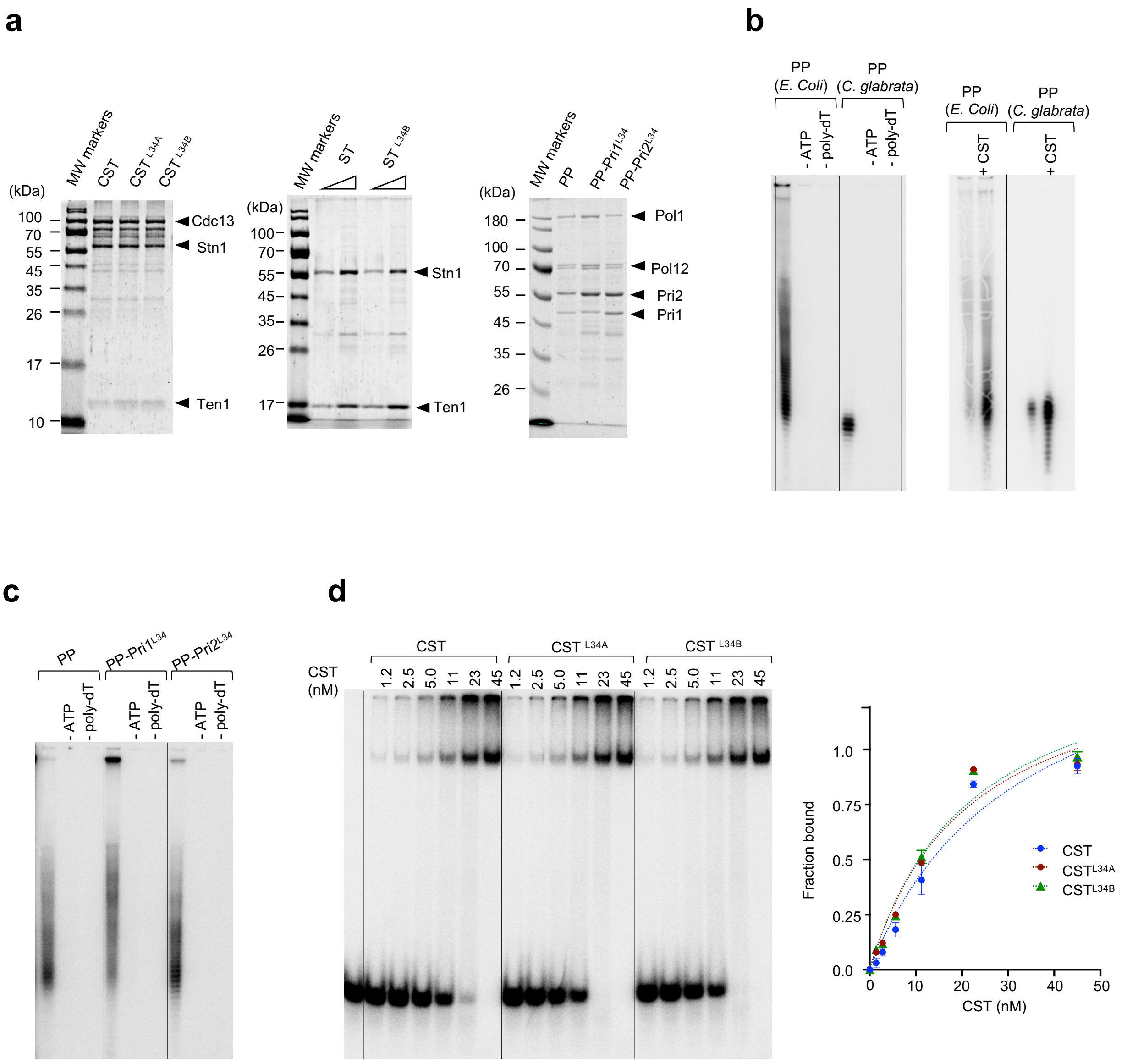

**Supp. Fig. 2. Characterizations of the CST and PP complexes utilized in this study**

**a.** The CST, ST and PP complexes containing wild type or mutated subunits were analyzed by SDS-PAGE and Coomassie staining.

**b.** **(Left)** The PP complexes purified from *E. coli* and *C. glabrata* were tested in parallel with regard to the dependence of product synthesis on the poly-dT template and ATP. Note that the PP complexes from *E. coli* synthesized a higher levels of longer products. This is most likely due to the excess of primase subunits in the preparation, which resulted in the synthesis of multimers of unit primer (~7-10 nt) before extension by DNA polymerase. **(Right)** The PP complexes purified from *E. coli* and *C. glabrata* were tested in parallel with regard to the stimulation of product synthesis by CST.

**c.** The PP complexes containing wild type subunits and mutated Pri1 or Pri2 proteins were tested in parallel with regard to the dependence of product synthesis on the poly-dT template and ATP.

**d.** The telomere DNA binding activity of CST, $CST^{L34A}$ and $CST^{L34B}$ were analyzed by EMSA using $P^{32}$-labeled CgTEL Gx1.5 as the substrate. Representative titration series are shown on the left and the quantified results plotted on the right.

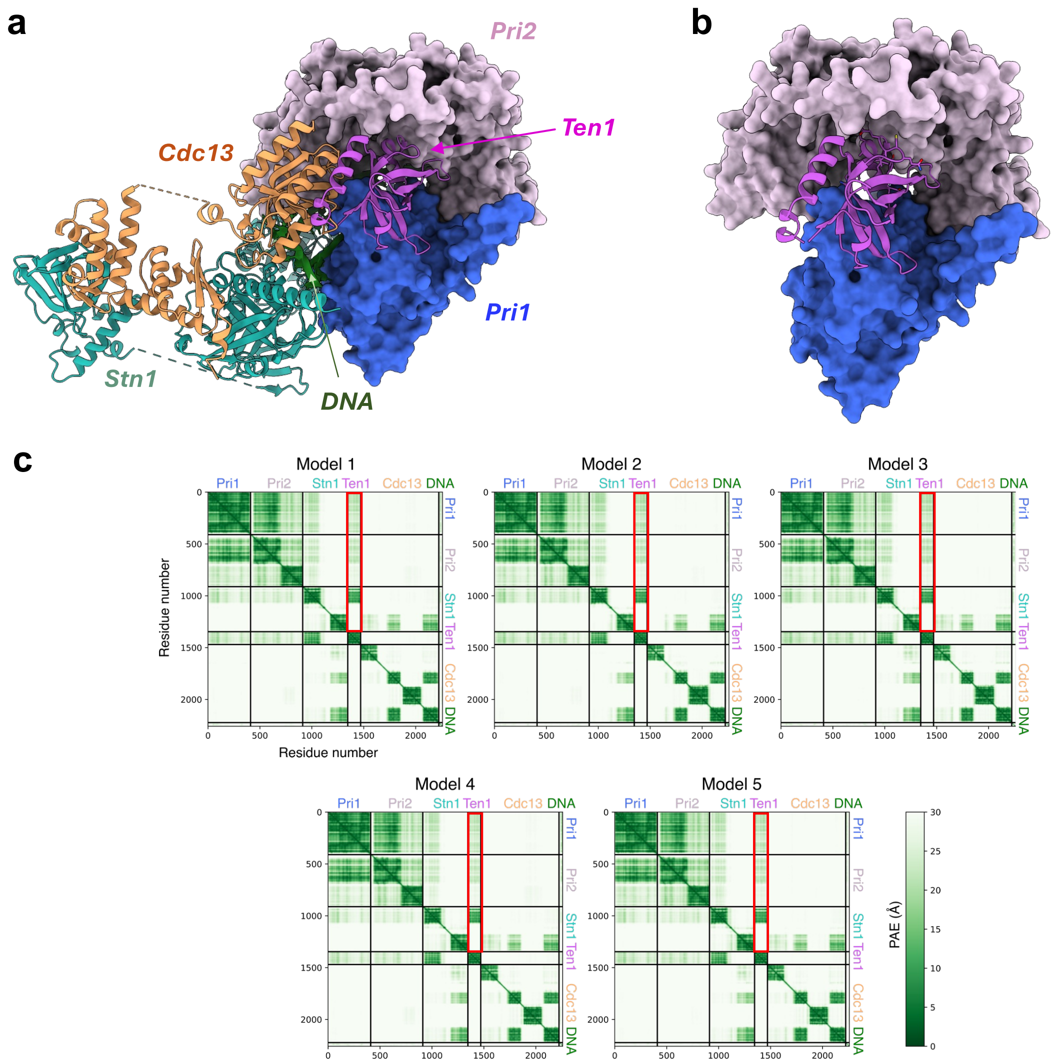

**Supp. Fig. 3. AlphaFold derived models of the *C. glabrata* CST-primase-DNA complex**

**a.** The top ranked model predicted by AlphaFold 3 using the *C. glabrata* Cdc13, Stn1, Ten1, Pri1 and Pri2 proteins as well as a 36-mer G-strand ssDNA ([TGTGGGGTCTGGGTGC]$_2$TGTG) is displayed. The CST subunits are shown in the designated colors using ribbon, and the primase subunits are shown in surface representations.

**b.** The Ten1, Pri1, and Pri2 proteins alone from the AlphaFold 3 model are displayed to illustrate the positioning of the Ten1 L34 loop adjacent to the Pri1/Pri2 protein-protein interface.

**c.** The expected position errors for the CST-primase-DNA models generated for all 5 models are displayed. The PAE predicted for Ten1 in relation to Pri1, Pri2 and Stn1 are highlighted with red rectangles.

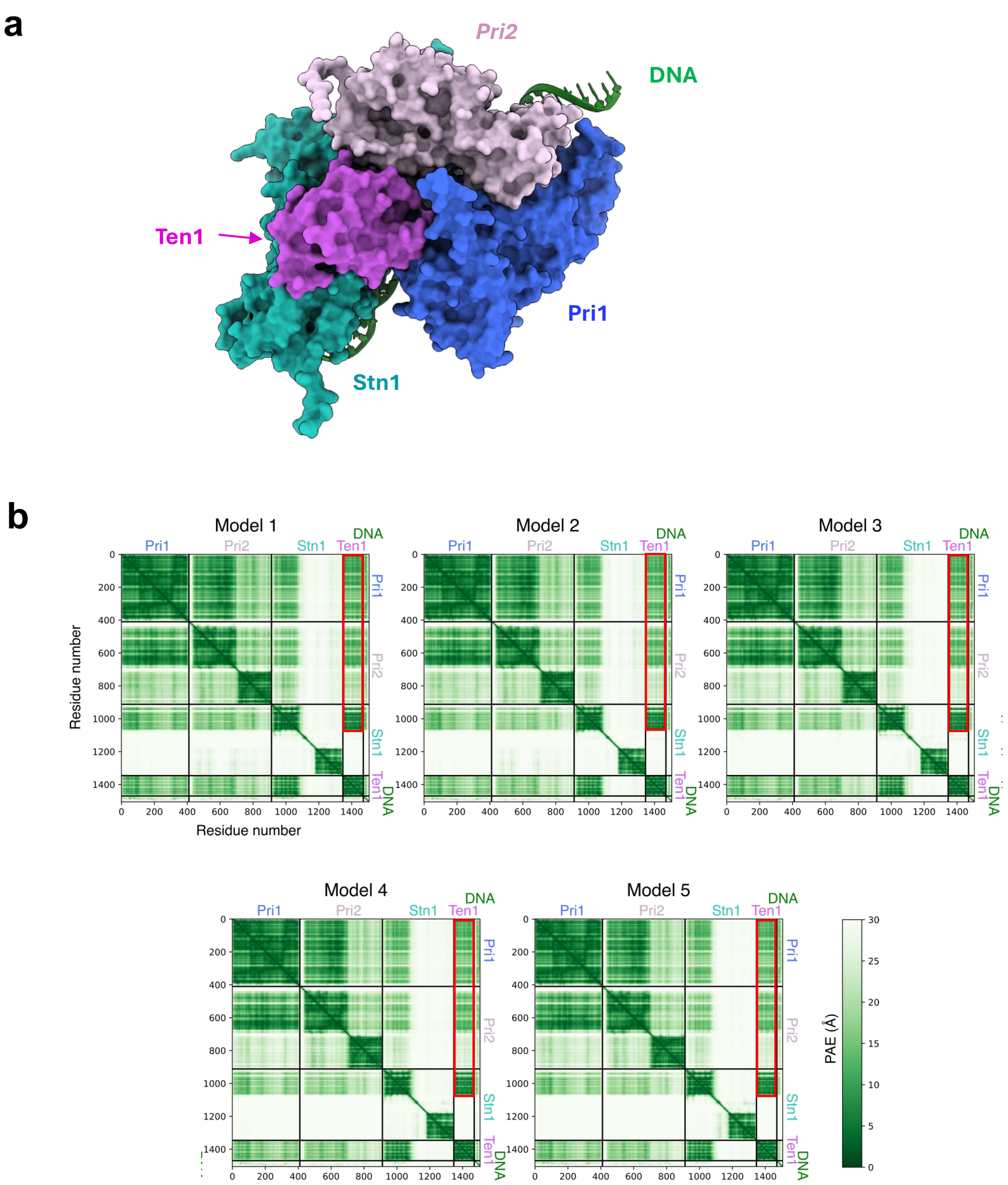

**Supp. Fig. 4. AlphaFold prediction for the *C. glabrata* ST-primase-DNA complexes**

**a.** The top ranked model for the *Cg*ST-primase-DNA complex derived from AlphaFold 3 using the specified proteins and a 36-mer C. *glabrata* telomere G-strand are displayed in surface representations.

**b.** The expected position errors for all 5 models of the *Cg*ST-primase-DNA models obtained from AlphaFold 3 are displayed. The PAE predicted for Ten1 in relation to Pri1, Pri2 and Stn1 are highlighted with red rectangles.

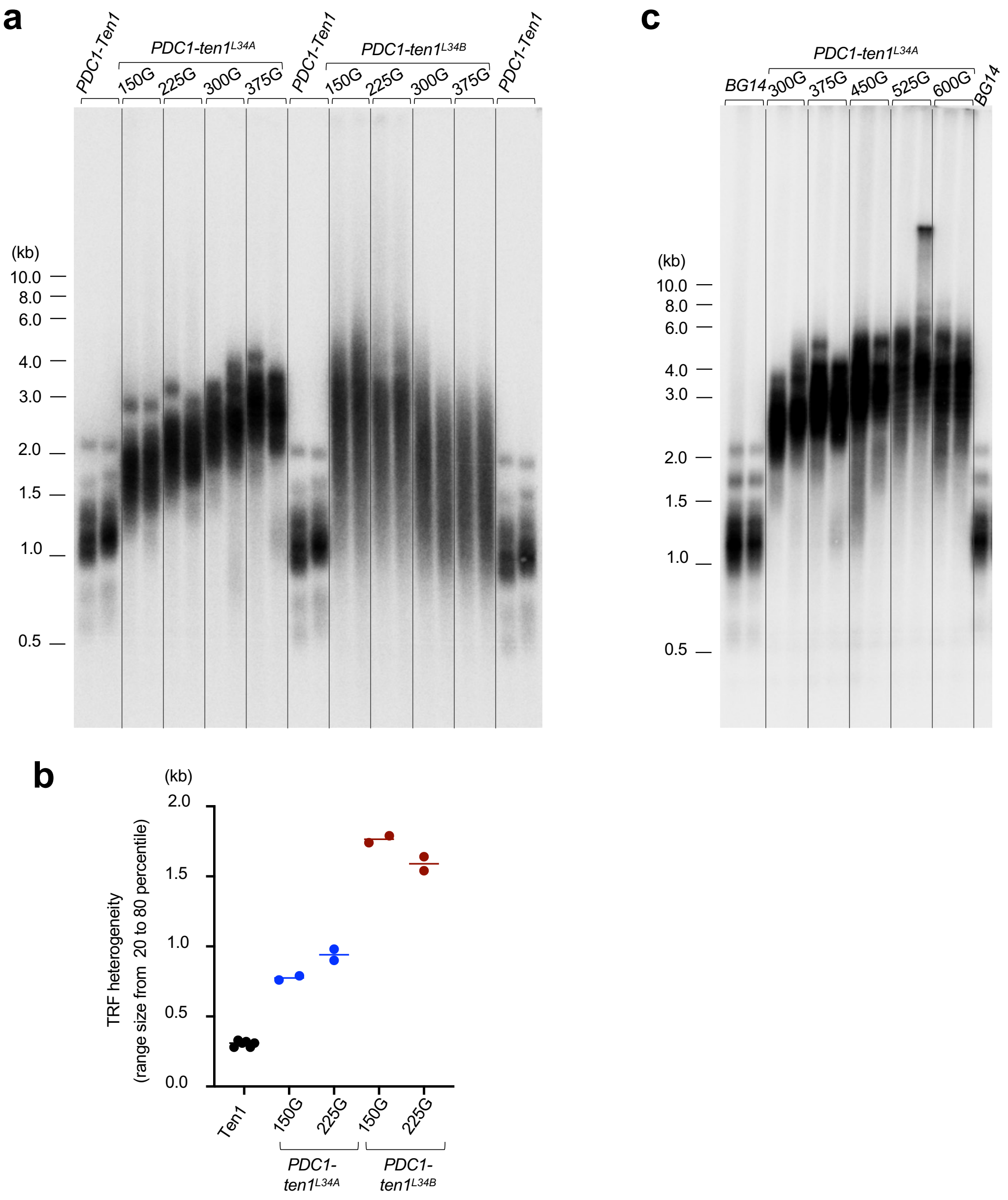

**Supp. Fig. 5. Telomere dynamics and heterogeneity induced by *ten1*$^{L34A}$ and *ten1*$^{L34B}$ during passage**

**a.** Genomic DNAs were isolated from the indicated strains that had been passaged for the designated number of generations, digested with *Eco*RI *and Alu*I, and subjected to TRF Southern analysis.

**b.** The heterogeneity of TRF lengths in the specified samples was determined based on the size difference between the $20^{th}$ percentile and the $80^{th}$ percentile TRF signals and plotted. For example, the $20^{th}$ and $80^{th}$ percentile TRFs for cells expressing *PDC1-Ten1* were about 0.9 and 1.2 kb, resulting in a heterogeneity estimate of about 0.3 kb.

**c.** Genomic DNAs were isolated from the *PDC1-ten1*$^{L34A}$ strain that had been passaged for the designated number of generations, digested with *Eco*RI *and Alu*I, and subjected to TRF Southern analysis.

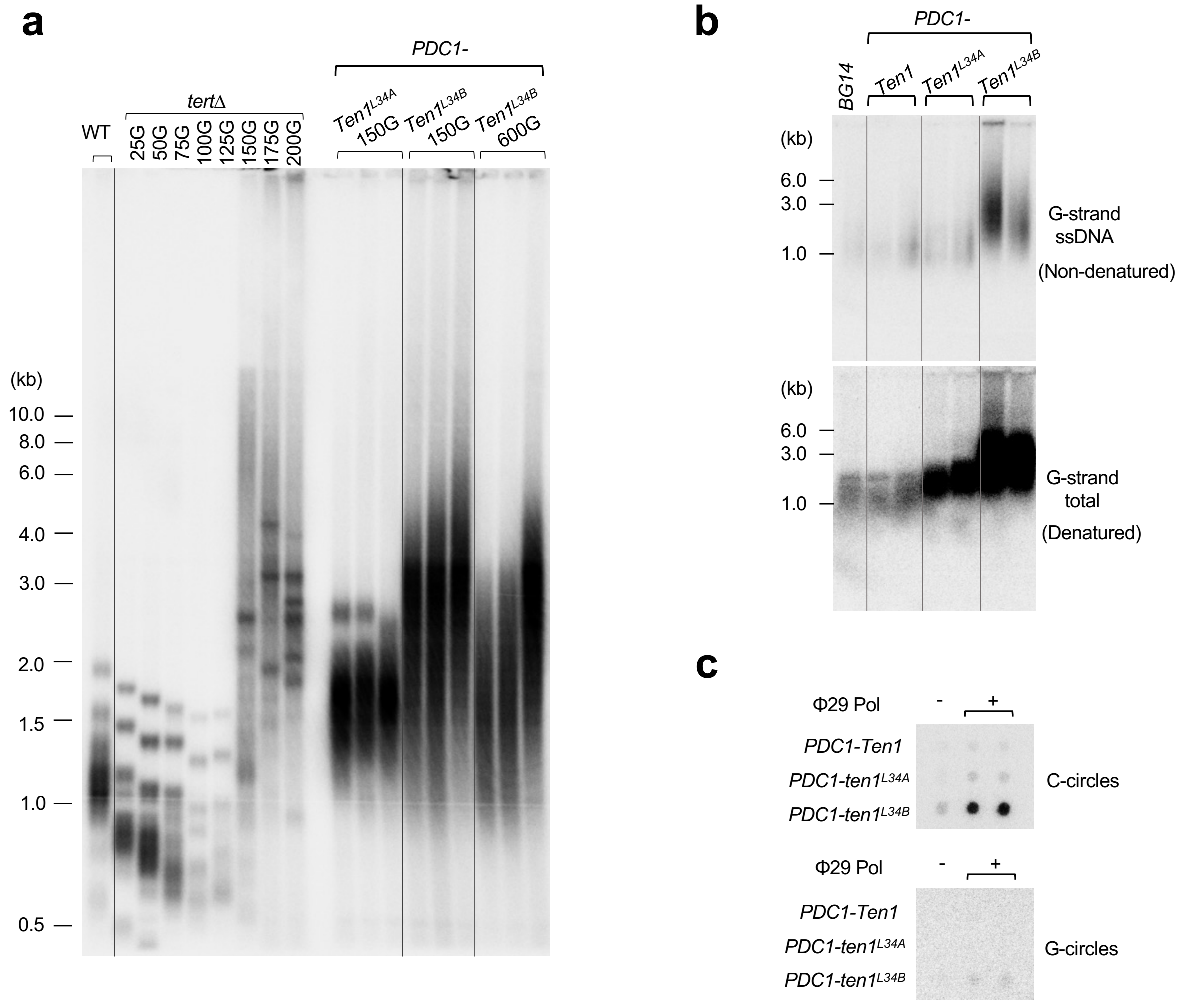

**Supp. Fig. 6. The *ten1*$^{L34B}$ strain exhibits hallmarks of telomere deprotection.**

**a.** Genomic DNAs were isolated from the indicated strains that had been passaged for the designated numbers of generations, digest with *Eco*RI and *Alu*I, and subjected to TRF Southern analysis.

**b.** Genomic DNAs were isolated from the indicated strains that had been grown for 75 generations, digest with *Eco*RI and *Alu*I, and subjected to in-gel hybridization analysis to detect G-strand ssDNA (top panel). The same gel was then denatured and rehybridized with the same probe to detect total G-strand DNA (bottom panel).

**c.** Genomic DNAs were isolated from the indicated strains that had been grown for 75 generations, digest with *Eco*RI, and subjected to C-circle and G-circle analysis.

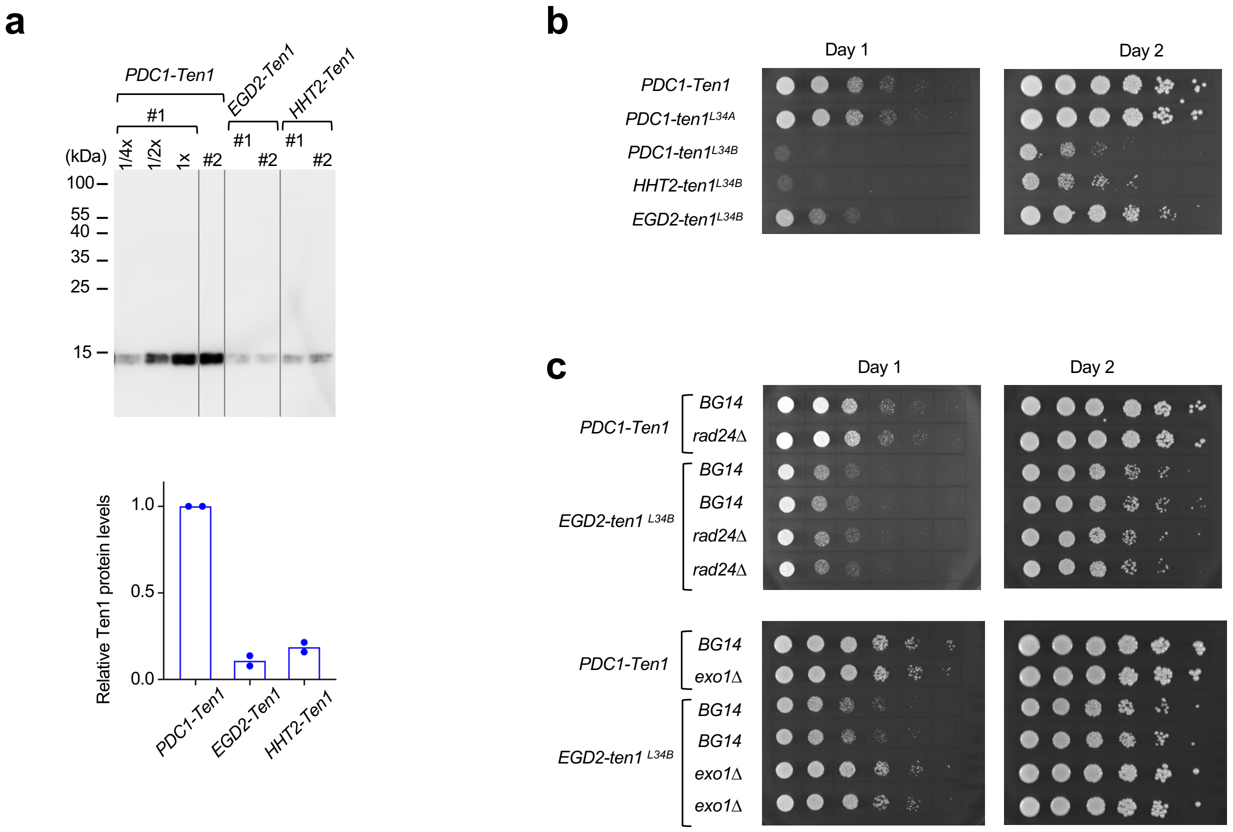

**Supp. Fig. 7. Comparisons of Ten1 expressed from the *PDC1*, *HHT2* and *EGD2* promoters; effects of *rad24*Δ and *exo1*Δ on the growth of cells expressing *EGD2-ten1*$^{L34B}$**

**a.** Western analysis of $FG_3$-tagged Ten1 in lysates from clones transformed with the indicated expression plasmids. Duplicated assay results are shown at the top and the quantifications plotted on the bottom. Note that a dilution series was performed using the one of the clones carrying *PDC1-Ten1* to generate a standard curve for quantitation (1$^{st}$ to 3$^{rd}$ lanes).

**b** and **c.** Serial dilutions of the indicated strains were spotted onto SD-ura semi-solid media and incubated at 30 degree. Images of the plates after 1 and 2 days of incubation are displayed.

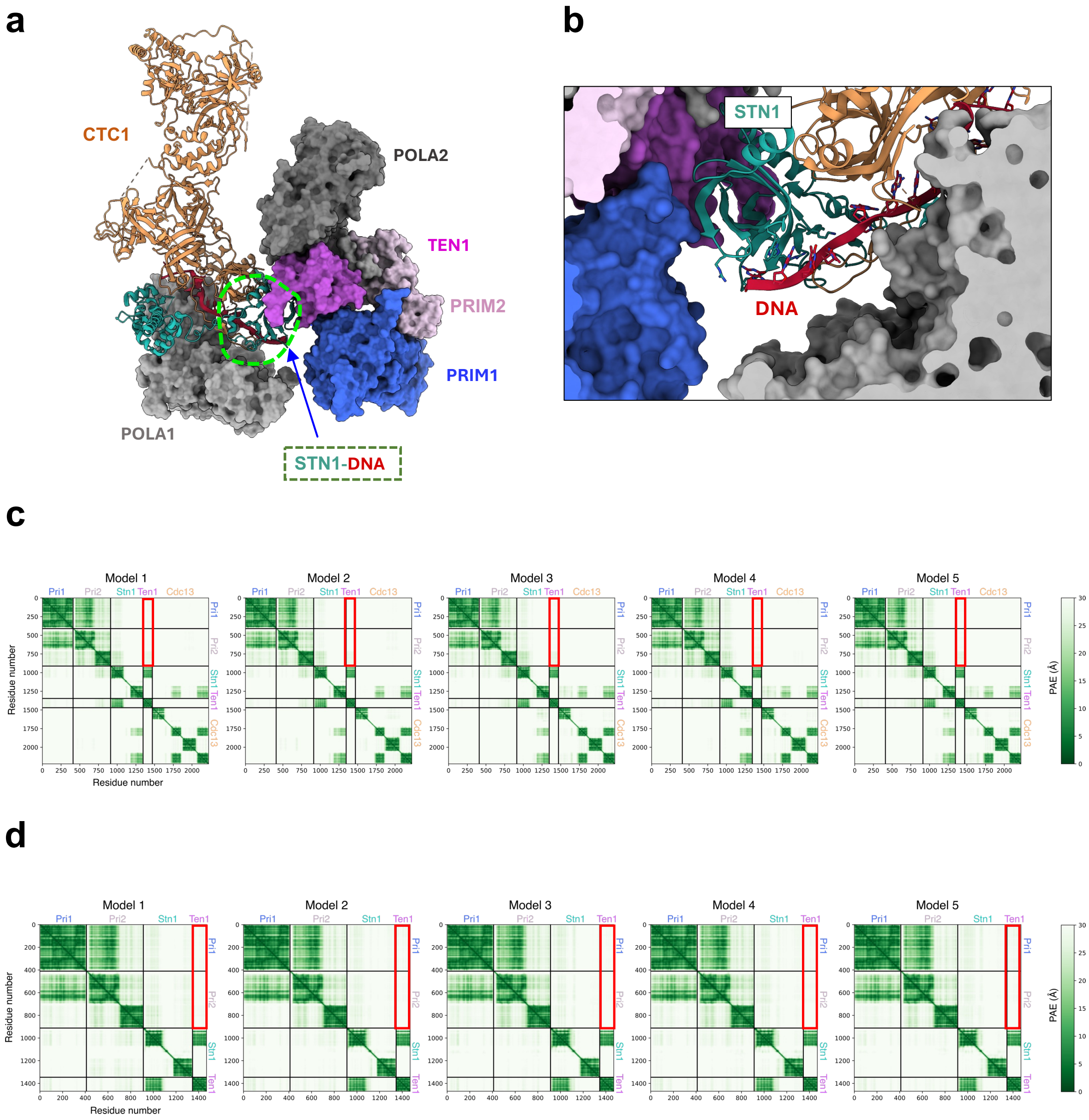

**Supp. Fig. 8. The role of Stn1-DNA interaction in the formation of the CST-PP-DNA PIC complex**

**a.** Structure of the human CST-PP-DNA pre-initiation complex illustrating the Stn1-DNA interaction (marked by a dashed circle) adjacent to the TEN1-PRIM1-PRIM2 interaction.

**b.** A more detailed view of the Stn1-DNA interaction in the human CST-PP-DNA pre-initiation complex is presented.

**c.** The PAE matrices for all 5 models of the *Cg*CST-primase complex (without DNA) generated by AF3 are displayed. The PAE predicted for Ten1 in relation to Pri1 and Pri2 are highlighted with red rectangles.

**d.** The PAE matrices for all 5 models of the *Cg*ST-primase complex (without DNA) generated by AF3 are displayed. The PAE predicted for Ten1 in relation to Pri1 and Pri2 are highlighted with red rectangles.

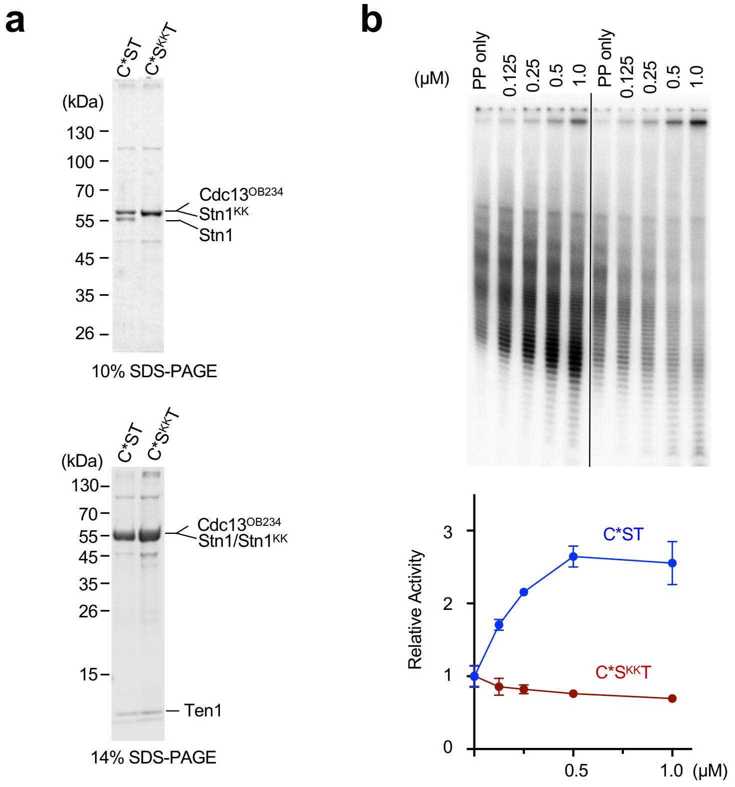

**Supp. Fig. 9. Disrupting Stn1-DNA interaction abrogated the stimulatory effect of *Cg*CST on PP activity *in vitro***

**a.** The C*ST and C*S$^{KK}$T complexes used in the PP assays are analyzed using different percentages of SDS-PAGE and Coomassie staining. Note that the Ten1 subunit has migrated off the bottom of the 10% gel owing to its small size. Also note that Stn1$^{KK}$ has reduced mobility relative Stn1 in SDS-PAGE and co-migrates with Cdc13$^{OB234}$.

**b.** The activity of the PP complex in synthesizing poly-rA-poly-dA from the poly-dT template was analyzed in the presence of increasing concentrations of C*ST and C*S$^{KK}$T. A representative series of assays are shown on top and the quantifications from 3 independent experiments plotted on the bottom.

**Supplementary Table 1. *C. glabrata* strains used in this study**

| **Alias** | ***Relevant Genotype*** | **Parental Strain** | **Reference** |
|---|---|---|---|
| *BG14* | *ura3Δ::Tn903 G418$^R$* | BG2 | Cormack and Falkow, 1999 |
| *exo1Δ* | *exo1Δ*::NAT$^R$(pBV65) | BG14 | This work |
| *rad24Δ* | *rad24Δ*::NAT$^R$(pBV65) | BG14 | This work |
| *rad52Δ* | *rad52Δ*::NAT$^R$(pBV65) | BG14 | This work |
| *tertΔ* | *tertΔ*::NAT$^R$(pBV65) | BG14 | This work |
| *LT* [a] | *pCU-PDC1-ten1$^{L34A}$* 225G | BG14 | This work |
| *LT/tertΔ* [b] | *pCU-PDC1-ten1$^{L34A}$* 225G, *tertΔ*::NAT$^R$(pBV65) | BG14 | This work |

[a] The BG14 strain carrying *pCU-PDC1-ten1$^{L34A}$* was passaged for ~ 225 generations to induce telomere lengthening before analysis or further manipulation.

[b] The *LT* strain (i.e., with elongated telomeres and carrying *pCU-PDC1-ten1$^{L34A}$*) was subjected to transformation with a *TERT* disruption cassette.

**Supplementary Table 2. Oligos used in this study**

| Name | Sequence 5' to 3' |
|---|---|
| **C. glabrata expression vectors** | |
| CgTen1-up-Bam | AATGGATCCATGAGCAAACTGGTGGTCG |
| CgTen1-dn-Not | AAGCGGCCGCACCGAGCAATTCACGGTC |
| CgTen1-up-Xba | AAGTCTAGAATGAGCAAACTGGTGGTCG |
| CgStn1-up-Xba | TAATCTAGAATGGAACATGGCGAGC |
| CgStn1-dn-NotI | AAGCGGCCGCATTATTGCATGTTTGAGTTG |
| **C. glabrata disruption cassette** | |
| Rad24-Aat-5UTR-F | cccgaaaagtgccacctgacgtcCTCCTCCAAAATCCTAG |
| Rad24-Pvu-5UTR-R | cctgcagcgtacgaagcttcagctgTTTGGTCCACTAGCCA |
| Rad24-SacII-3UTR-F | ctagtggcctatgcggccgcggTGACATGATGGTTTCCA |
| Rad24-HpaI-3UTR-R | ggccgattcattaatgcaggttaacCTTGAAGGAAGTGGTGC |
| Rad52-Aat-5UTR-F | cccgaaaagtgccacctgacgtcTGAGCTGTCTCCTTTGGTC |
| Rad52-Pvu-5UTR-R | cctgcagcgtacgaagcttcagctgCGCTTAAGGTTAAGTCACC |
| Rad52-SacII-3UTR-F | ctagtggcctatgcggccgcggCACCATGATCCATGGATTG |
| Rad52-HpaI-3UTR-R | ggccgattcattaatgcaggttaacCTTCATTCTTCGGTCATCATC |
| Exo1-Aat-5UTR-F | cccgaaaagtgccacctgacgtcCTTATTCTCTCCAGCAGC |
| Exo1-Pvu-5UTR-R | cctgcagcgtacgaagcttcagctgTCAATGACCTCTAGAACTG |
| Exo1-SacII-3UTR-F | ctagtggcctatgcggccgcggCTACATATAACCAGTCTGC |
| Exo1-HpaI-3UTR-R | ggccgattcattaatgcaggttaacGATAGTCGGCACAAATTC |
| CgTERT-5UTR-F-Hpa | TCCGTTAACGCCAATACTTCTCTGACCATTC |
| CgTERT-5UTR-R-SacII | TTACCGCGGTCATTTATTATCTGTTAGCATAGTA |
| CgTERT-3UTR-F-BsiWI | TTACGTACGCGGAATATCTAGAAATATGAACTCG |
| CgTERT-3UTR-R-AatII | TTTGACGTCCAGAAAGGATGTTGCTGAGA |
| **Mutagenesis** | |
| CgTen1-L12-F | GAGCTGGTCGCGGCTGCTGCTGCGGCTGCAGCTATCATAGTGCTCAG AAATGCGCCT |
| CgTen1-L12-R | CACTATGATAGCTGCAGCCGCAGCAGCAGCCGCGACCAGCTCTTCCA CTCGCCCTAT |
| CgTen1-L23-F | GTGCTCAGAGCTGCGGCTGCAGCAGCGGCTGCTGCAGGTAAGTTGCG AGTTTCTGTTGGA |
| CgTen1-L23-R | CAACTTACCTGCAGCAGCCGCTGCTGCAGCCGCAGCTCTGAGCACTA TGATACTGATATA |
| CgTen1-L34A-F | GTTTCTGTTGCAGCTGCGGCAGCTGCCGCATTAGCCACCGCTATTGGA GAGCTA |
| CgTen1-L34A-R | GGTGGCTAATGCGGCAGCTGCCGCAGCTGCAACAGAAACTCGCAACT TACCGTC |
| CgTen1-L34B-F | AGATTAGCCGCCGCTGCTGCAGCAGCAGCCGCAGGGAGTGCTGTAGA CGCTATGATT |
| CgTen1-L34B-R | AGCACTCCCTGCGGCTGCTGCTGCAGCAGCGGCGGCTAATCTGGAAT AGCACGCATC |
| CgTen1-L5C-F | TTGCTGGACGCGGCAGCTGCATCAGTGGGTGAAGTTGAAGCACTG |

| | |
|---|---|
| CgTen1-L5C-R | ACCCACTGATGCAGCTGCCGCGTCCAGCAACTCATATGCTACTTC |
| CgPri1-L34$^{Tar}$-F | GATTCGCTGGCGGCAGCTGCGGCTGCACCATTAGAGAAGGAACTTGT ATTC |
| CgPri1-L34$^{Tar}$-R | CTCTAATGGTGCAGCCGCAGCTGCCGCCAGCGAATCACGCTCCTTTG GTGG |
| CgPri2-L34$^{Tar}$-F | CTAATTAAAGCAGCCGCAGCCGCAGCAGCATTGAACGAGGACGACAG ACTGATA |
| CgPri2-L34$^{Tar}$-R | CTCGTTCAATGCTGCTGCGGCTGCGGCTGCTTTAATTAGTTCTTCTTGT AGTTT |
| CgStn1-KK69_74EE-F | ATTGGTGTCGAGTATATGTGGCTCGAGGGAGATGACTACGCCGTAATT |
| CgStn1-KK69_74EE-R | GTCATCTCCCTCGAGCCACATATACTCGACACCAATCACTGTACCAAC |
| CgStn1-F127E-F | AATACCGAAGAGAATGAGCTGGAAGTAGGTTTTATC |
| CgStn1-F127E-R | CAGCTCATTCTCTTCGGTATTGTAAACACCACATAC |
| **hybridization, EMSA assays** | |
| CgTEL-G1.5 | TGTGGGGTCTGGGTGC TGTGGGGT |
| CgTEL-C1.5 | ACCCCACA GCACCCAGACCCCACA |
| CgTEL-G3 | $[TGTGGGGTCTGGGTGC]_3$ |
| CgTEL-C3 | $[GCACCCAGACCCCACA]_3$ |
